# Enhance genome editing efficiency and specificity by a quick CRISPR/Cas9 system

**DOI:** 10.1101/708404

**Authors:** Yingying Hu, Zhou Luo, Jing Li, Dan Wang, Hai-Xi Sun, An Xiao, Da Liu, Zhenchao Cheng, Jun Wu, Yue Shen, Xun Xu, Bo Zhang, Jian Wang, Ying Gu, Huanming Yang

## Abstract

CRISPR/Cas9 is a powerful genome editing tool that has been successfully applied to a variety of species, including zebrafish. However, targeting efficiencies vary greatly at different genomic loci, the underlying causes of which were still elusive. Here we report a quick CRISPR/Cas9 system, designated as qCas9, which exhibits accelerated turnover of Cas9 protein in zebrafish. Our data showed that qCas9 significantly improved targeting efficiency, including both knock-out and knock-in in F_0_ embryos, and yielded higher germline transmission rate in founder screen. Importantly, qCas9 showed little to no off-target editing in zebrafish and profoundly reduced off-target effect in HEK293T cell line. In summary, our findings demonstrate that qCas9 is a simple, economic and highly effective method to improve genome editing efficiency in zebrafish embryos and also holds great potential in reducing off-target effect in mammalian cell lines.

## Introduction

Natural CRISPR systems found in eubacteria and archaea, including type I, II and III systems, use small RNA and CRISPR-associated protein (Cas) to target invading foreign DNAs^1-3^. Type II CRISPR/Cas system that relies on the endonuclease Cas9 from *Streptococcus pyogenes* was found to induce RNA-guided site-specific DNA-cleavage in vitro^4^, in cultured cell lines^5,6^, and in vivo, e.g. mouse^7,8^, zebrafish^9,10^, fruit fly^11^ and *C*.*elegans*^12^. After being introduced into a cell, the Cas9 protein forms a complex with a synthetic RNA molecule containing a guide sequence complementary to a specific DNA sequence (protospacer), and thereby enabling the introduction of a double strand break (DSB) in the genomic locus of interest. The DSB is mainly repaired by two cellular DNA damage repair pathways, the error-prone non-homologous end joining (NHEJ) pathway^13^ or error-free homology-directed repair (HDR) pathway^14^.

Zebrafish is an important vertebrate model organism for functional genomics study and is readily amenable to genome editing via ZFN^15^, TALEN^16,17^ and the CRISPR/Cas9 system^10,18^. Despite the ease and versatility of the CRISPR/Cas9 system, however, mutagenesis efficiency of each Cas9/gRNA target site varies greatly ranging from almost 0% to nearly 100%. Although many online programs are available for predicting gRNA efficiency ^19-21^, the majority of these prediction rules were derived from data of cultured cells and thus of less predicative value for gRNA design in zebrafish^22^. Low mutagenesis efficiency requires a large amount of labor and time to obtain a desired zebrafish mutant line. One obstacle for achieving efficient genome targeting in zebrafish is the rapid cell division after fertilization, leaving limited time window for genome editing tools to function; therefore leading to high mosaicisms of edited zebrafish embryos and low germline transmission rate. To overcome this, several methods have been developed. One way to improve genome targeting efficiency in zebrafish is to bypass mRNA translation through the use of Cas9 protein, which resulted in rapid and higher targeting efficiency^23^. Another strategy involves direct injection of Cas9/gRNA into oocytes rather than zygotes^24^. In addition, the use of species-specific codon optimized Cas9^25,26^ and the addition of non-homologous single-stranded DNA^27^ were also shown effective. Despite these advances, however, new strategies to increase targeting efficiency in zebrafish, especially with easy manipulation and low cost are still desirable.

The concerns of off-targeting effect also hinder CRISPR/Cas9’s applications in agriculture and medicine^28^. One key to minimize off-target editing is to increase on-target specificity. Many efforts have been devoted to the optimization of CRISPR/Cas9 system with high specificity, such as the use of truncated guide RNA^29^, Cas9 nickase^30^, FokI-dCas9^31^, structure guided engineering of Cas9^32,33^, using small molecules or light to temporally manipulate Cas9 activity^34-37^ and partial replacement of RNA nucleotides with DNA nucleotides in crRNA^38^.

Another important consideration to reduce off-targeting effect is shortening the duration time of Cas9 protein. To the end, we hypothesize that by introducing Cas9 protein earlier and decreasing its half-life, both higher on-target editing efficiency and lower off-target effect can be achieved.

Proteins have variable turnover rates and half-lives. It was proposed that most of the short lived protein contained a region enriched with proline (P), glutamic acid (E), serine (S), and threonine (T), which was called the PEST sequence or PEST domain^39^. Mouse ornithine decarboxylase, structural 1 (ODC1), for example, is one of the short lived proteins that contains the PEST sequence at its C terminus^40^. PEST sequence acts as a proteolytic signal for rapid protein degradation by ubiquitin-26S proteasome^41,42^, and thereby shortening protein half-life^43,44^. This feature of the PEST sequence has been harnessed for the generation of a EGFP-PEST transcription reporter, which enabled a more sensitive way to monitor dynamic biological signals than that of conventional EGFP reporter^45^. It is also important to note that PEST sequence not only facilitated rapid degradation of EGFP-PEST fusion protein as expected, but also accelerated the translation of EGFP-PEST fusion protein in zebrafish, therefore holding the potential to achieve faster turnover rate of protein of interest.

In this study, we fused PEST domain to Cas9 and named this fusion protein as quick Cas9 (qCas9) due to the rapid Cas9 protein turnover rate enabled by the PEST sequence. Strikingly, qCas9 greatly enhanced on-target efficiency in zebrafish embryos, and this effect was dosage dependent, worked at various time points and yielded higher germline transmission rate in founder screen. qCas9/*tyr* gRNA targeting led to higher ratio of embryos with severe loss-of-pigmentation phenotype than that of conventional Cas9/*tyr* gRNA targeting and also resulted in higher targeted knock-in efficiency. Importantly, the enhancement of on-target editing by qCas9 did not increase off-target editing when compared to Cas9. In addition, human-codon-optimized Cas9-PEST (quick hCas9, qhCas9) significantly reduced off-target editing of VEGFA sites in HEK293T cell line, though in this case on-target editing efficiency was only marginally improved. In summary, here we develop a safer and more efficient CSISPR/Cas9 system by fusing PEST sequence to the C terminal of Cas9 protein. We envision this strategy can easily be applied to other species and other CRISPR based genome editing tools, like CRSIPR/Cpf1 and CRSIPR/Base editors.

## Results

### PEST sequence accelerated mRNA translation

To test and validate the function of PEST sequence in zebrafish, a pCS2-EGFP-PEST vector was constructed by fusing PEST sequence from mouse ODC1 (amino acid 422-461) to the C-terminal of EGFP. Both capped EGFP and EGFP-PEST mRNA were in vitro transcribed by SP6 RNA polymerase and injected into one-cell stage zebrafish embryos. As early as 1 hour post fertilization (hpf), green fluorescence could be observed in EGFP-PEST mRNA injected embryos. The fluorescence intensity gradually increased as development proceeded and then faded away after 24 hpf. In contrast, GFP signal could not be observed until 2 hpf in EGFP mRNA injected embryos. At 24 hpf, EGFP mRNA injected embryos still maintained high level of fluorescence intensity (**Figure 1**). Taken together, these results suggest that PEST sequence can accelerate both mRNA translation and protein degradation.

**Figure 1.**
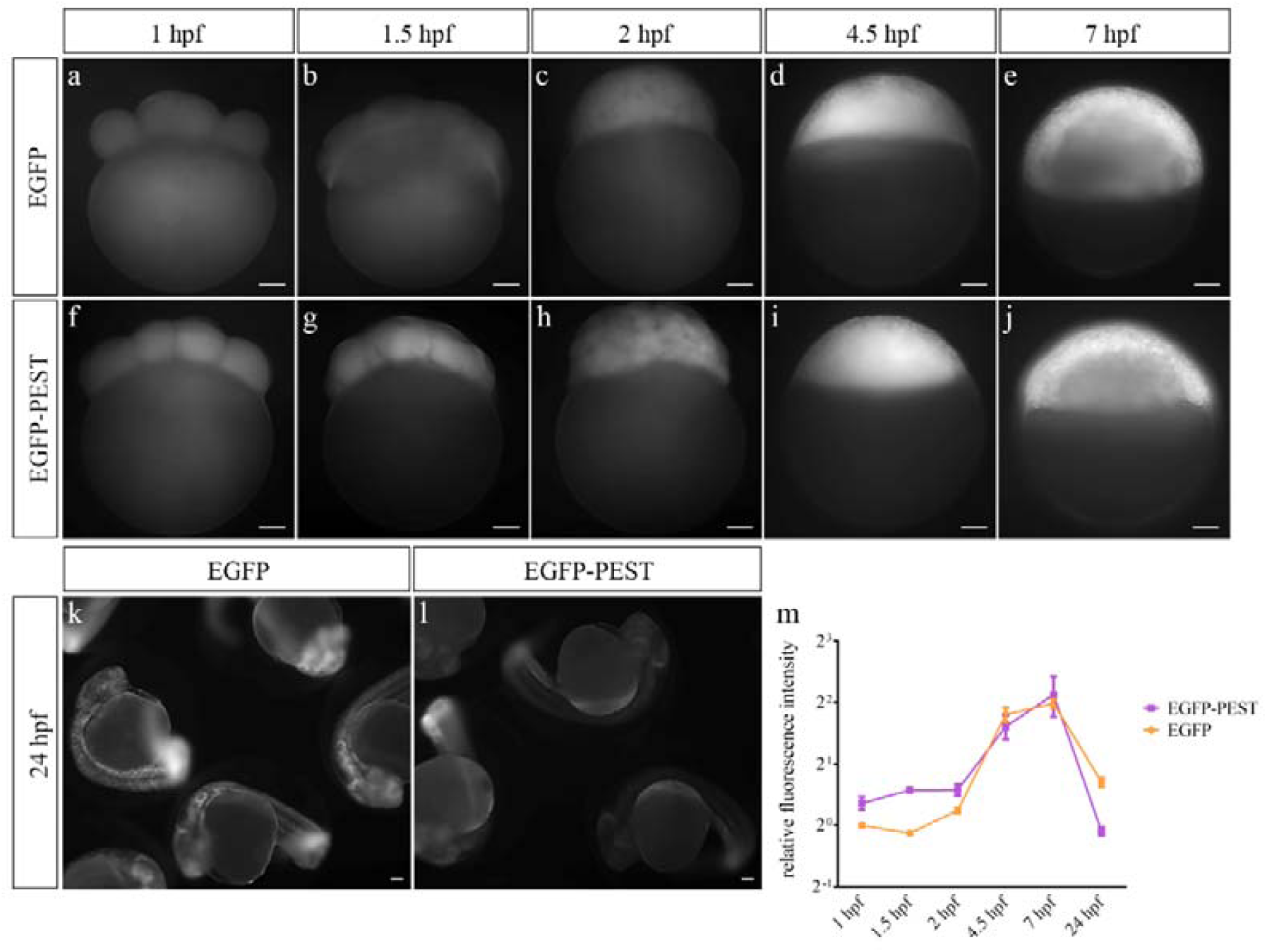
PEST sequence facilitated EGFP mRNA translation and degradation in zebrafish embryos. Fluorescence images of EGFP mRNA injected embryos (a, b, c, d, e, k) and EGFP-PEST mRNA injected embryos (f, g h, i, j, l) at 1 hpf, 1.5 hpf, 2 hpf, 4.5 hpf, 7 hpf and 24 hpf. Relative fluorescence intensity was calculated in (m). The results showed quicker expression of EGFP-PEST mRNA before 2 hpf and rapid degradation after 24 hpf.

### qCas9 outperformed Cas9 in on-targeting efficiency

The zebrafish-codon-optimized Cas9 (zCas9) flanked by both N and C terminal nuclear location signal (NLS) was used as previously described^25^ and served as the control Cas9 in this study. The PEST coding sequence (amino acid 422-461) (**Figure 2b**) from mouse *Odc1* was cloned and inserted to the C terminus of Cas9 to produce Cas9-PEST fusion protein (**Figure 2a**). Both Cas9 and Cas9-PEST mRNA were in vitro transcribed for embryo micro-injection. Considering the rapid protein turnover enabled by the PEST sequence, we named the Cas9-PEST fusion construct as the quick Cas9, qCas9.

**Figure 2.**
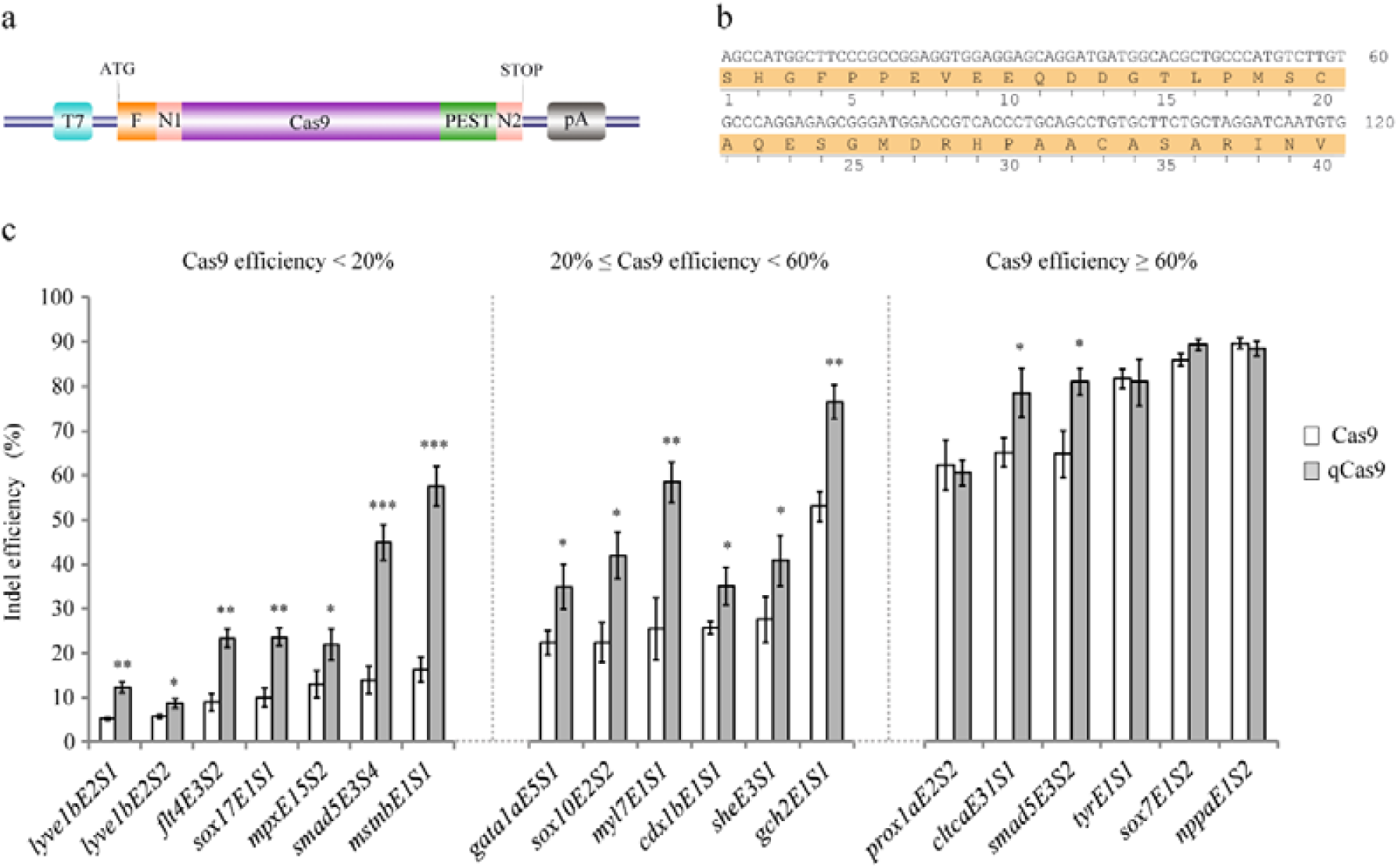
qCas9 improved targeting efficiency in multiple zebrafish endogenous sites. (a) Schematic design of Cas9-PEST construct. The zebrafish codon optimized Cas9 was flanked by 3x FLAG and SV40 nuclear localization signal (NLS) at 5’ end and nucleoplasmin NLS and polyA sequence at 3’ end. The PEST sequence from mouse *Odc1* was cloned and inserted into the C terminal of Cas9. T7 promoter was used for in vitro transcription of Cas9-PEST mRNA. (b) The PEST sequence: 120 bp nucleotide sequence of mouse *Odc1* PEST domain, the respective amino acid sequence (422aa-461aa) was listed below. (c) A mixture of 400pg Cas9 or Cas9-PEST mRNA and 50 pg guide RNA was co-injected into one-cell stage of zebrafish embryos. For each target site, genomic DNA of 18 embryos (3 embryos per group, 6 groups) were extracted and indel efficiencies were determined by restriction fragment length polymorphism (RFLP) assay or T7 Endonuclease I assay at 24 hpf. Two-tailed unpaired Student’s t-test was used for statistical analysis with *, **, and *** represent *p* value < 0.05, 0.01 and 0.001, respectively.

To test the efficacy and efficiency of qCas9 in zebrafish, 23 gRNAs targeting various genomic sites were designed and evaluated for targeting efficiency using restriction-endonuclease-resistant restriction fragment length polymorphism (RFLP) assay or T7 Endonuclease I assay (see Methods). Except for 4 inactive target sites, 19 gRNAs showed targeting efficiencies ranging from 5% to 89% using Cas9 mRNA. The 23 gRNAs were then divided into 4 groups (inactive, low, medium and high) according to targeting efficiency (0%, <20%, 20%-60% and >60%) and tested using the same dosage of qCas9 mRNA. The results showed that co-injection of qCas9 mRNA and gRNA greatly increased targeting efficiencies of 13 sites in low and medium groups, while had less impact on sites with high efficiencies (**Figure 2c**). Note that 1 of the 4 inactive sites showed weakly detectable efficiency about 1% using qCas9 and the result was confirmed by Sanger sequencing. The data demonstrates that qCas9 can significantly improve targeting efficiencies of sites with low and medium efficiencies, and has only marginal effects on inactive and highly active sites. The detailed target sites information could be found in **Supplementary Table 1** and target site efficiency comparison data could be found in **Supplementary Fig. 1**.

### qCas9 increased targeting efficiency at various time points

Next, we examined qCas9 activities at 7 time points during zebrafish development (1 hpf, 2 hpf, 3 hpf, 6 hpf, 10 hpf, 24 hpf and 48 hpf). The gRNA target sites for *smad5*E3S4, *gch2*E1S1 and *sox7*E1S2 were chosen from our on-targeting efficiency test experiment as representatives for low, medium and high targeting efficiency, respectively. No targeting efficiency was detected at 1 hpf and 2 hpf for both Cas9 and qCas9 mRNA injected embryos, indicating the lack of sufficient functional Cas9 protein before 2 hpf. For both Cas9 and qCas9 mRNA injected embryos, on-target editing was first detected at 3 hpf using RFLP assay, gradually increased, and then culminated at 24 hpf for *smad5*E3S4 and *sox7*E1S2 and at 10 hpf for *gch2*E1S1. Faster temporal dynamics and improved targeting efficiency were consistently observed in qCas9 versus Cas9 at all the time points examined for *smad5*E3S4 and *gch2*E1S1 (**Figure 3a and 3b**). Though no significant difference of *sox7*E1S2 targeting efficiency between Cas9 and qCas9 was observed at 24 hpf and 48 hpf, qCas9 showed better performance before 10 hpf than that of Cas9 (**Figure 3c**). Targeting efficiency for *smad5*E3S4, *gch2*E1S1 and *sox7*E1S2 at different time points could be found in **Supplementary Fig. 2**.

**Figure 3.**
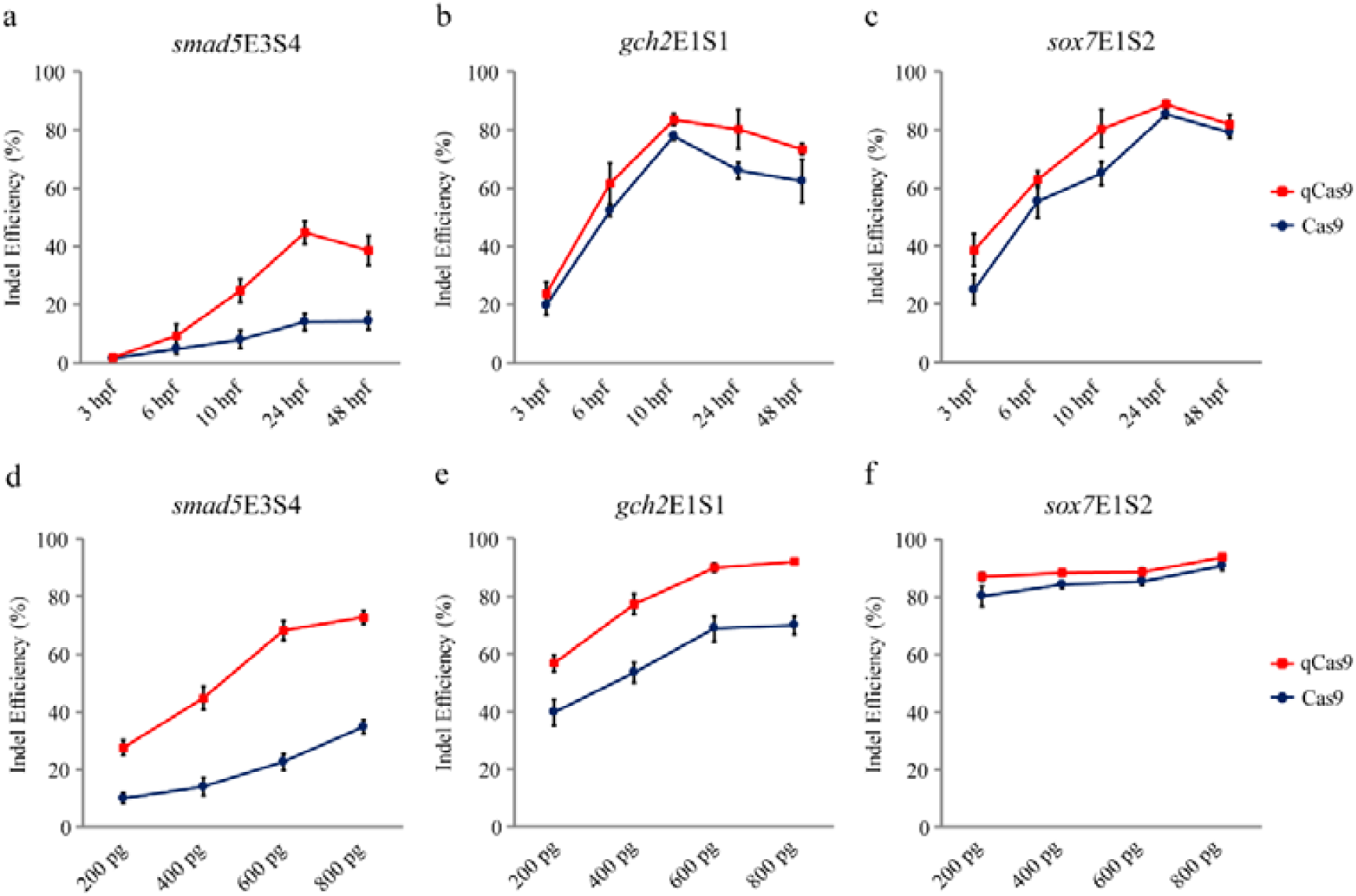
Time-course and dosage-dependent targeting efficiency of qCas9. (a-c) Dosage dependent experiment of Cas9 and qCas9. Different amount of Cas9 or qCas9 mRNA were co-injected with 50 pg guide RNA. The *smad5*E3S4 (a), *gch2*E1S1 (b) and *sox7*E1S2 (c) were chosen as representatives of low, median and high Cas9 indel efficiencies, respectively. (d-f) Time course experiment of Cas9 and qCas9. A mixture of 400pg Cas9 or qCas9 mRNA and 50 pg guide RNA was co-injected. Genomic DNA from 3 hpf, 6 hpf, 10 hpf, 24 hpf and 48 hpf embryos were extracted for indel efficiency analysis. The target sites *smad5*E3S4 (d), *gch2*E1S1 (e) and *sox7*E1S2 (f) were chosen as representatives of low, median and high Cas9 indel efficiencies, respectively. The results revealed that qCas9 outperformed Cas9 in all the situations examined.

### The enhancement of targeting efficiency was dosage-dependent

To test whether the improvement of targeting efficiency by qCas9 is dosage-dependent, different concentrations (200 pg, 400 pg, 600 pg and 800 pg) of Cas9 and qCas9 mRNA (**Supplementary Fig. 3**) were co-injected with 50 pg gRNA into one-cell stage embryos and the targeting efficiencies were determined at 24 hpf. Again, the gRNA target sites for *smad5*E3S4, *gch2*E1S1 and *sox7*E1S2 were used to represent low, medium and high targeting efficiency, respectively. In consistent with previous studies^25,26^, the targeting efficiencies for all 3 sites were dosage-dependent. Also note that qCas9 outperformed Cas9 in all dosages examined (**Figure 3d-f, Supplementary Fig. 4**).

### qCas9 generated more pronounced founder phenotype

To test whether qCas9 could facilitate phenotypic analysis in founder (F_0_) zebrafish, *tyrosinase* (*tyr*) was chose as the target gene since the loss-of-pigmentation phenotype of *tyr* mutant can be readily observed^46^. Co-injection of Cas9 mRNA and *tyr*E1S1 gRNA showed ∼80% targeting efficiency as previously reported^26^. The high targeting efficiency led to pigmentation defects in F_0_ embryos at 60 hpf (**Figure 4a**). Although qCas9 had marginal effects over Cas9 on highly efficient sites such as the *tyr*, interestingly, we observed that the percentage of embryos with strong pigmentation phenotype (lost more than 80% pigmented melanophores) was increased from 69.0% to 79.3% (**Figure 4b**). These results demonstrate that qCas9 could accentuate melanin pigmentation phenotype even when targeting efficiency was not enhanced.

**Figure 4.**
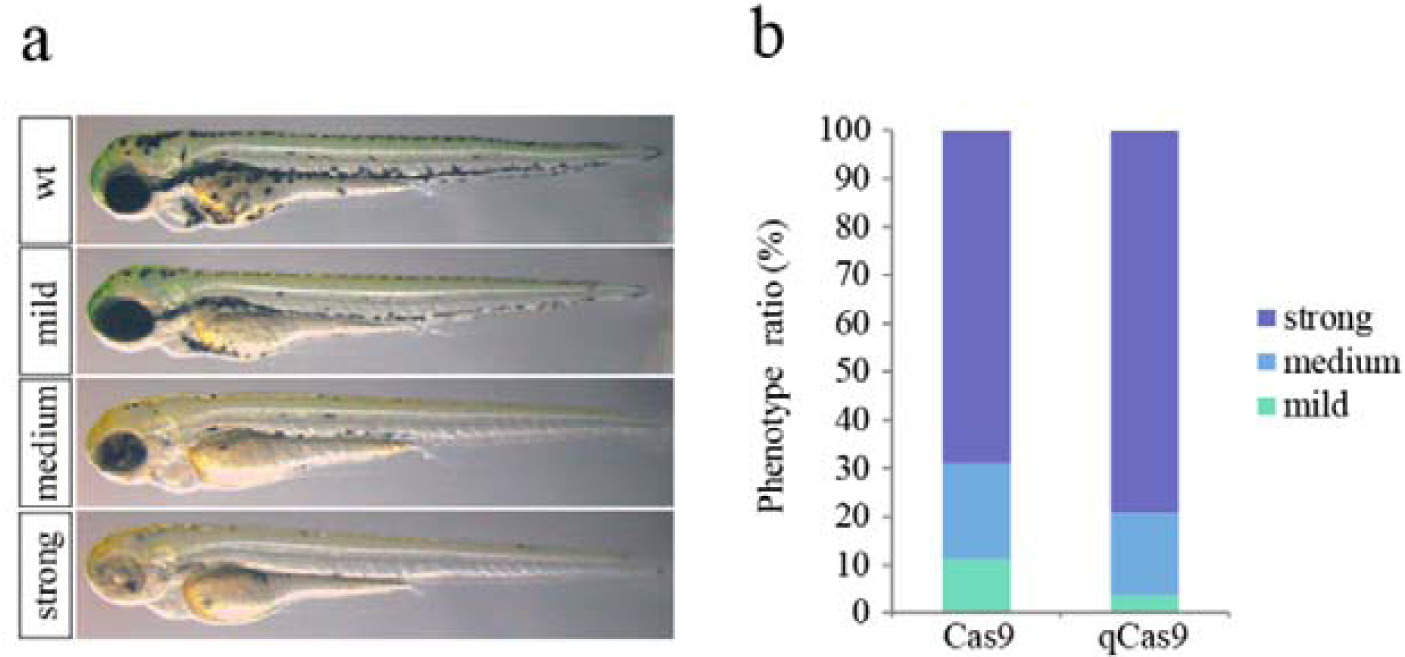
Disruption of *tyr* by qCas9 led to more embryos with pigmentation defects in F_0_ generation. (a) Co-injection of 400 pg Cas9 or qCas9 mRNA and 50 pg *tyr*E1S1 guide RNA led to high indel efficiency and induced loss-of-pigmentation phenotype in founder embryos at 60 hpf. (b) Ratio of embryos with different degree of pigmentation defects. Mild, medium and strong phenotype were correlated to embryos with >80% pigmented melanophores, 20%-80% pigmented melanophores and <20% pigmented melanophores, respectively. Though the indel efficiency was not significantly improved by qCas9, the ratio of embryos with strong pigmentation phenotype was increased from 69.0% to 79.3%.

To explore whether Knock-In (KI) efficiency could also be improved by qCas9, two donor plasmids were constructed for KI to *acta1a* and *nppa* loci (**Figure 5a**). After co-injection of KI donor plasmids, *acta1a*E1S1 gRNA and qCas9 or Cas9 mRNA, 18.7% of embryos from qCas9 group showed EGFP fluorescence in muscle at 24 hpf while only 5.9% of embryos from Cas9 group were EGFP positive. For the *nppa*E1S2 locus, 7.8% of embryos from qCas9 group showed tdTomato fluorescence in heart at 24 hpf while only 4.7% of embryos from Cas9 group were tdTomato positive (**Figure 5b** and **5c**).

**Figure 5.**
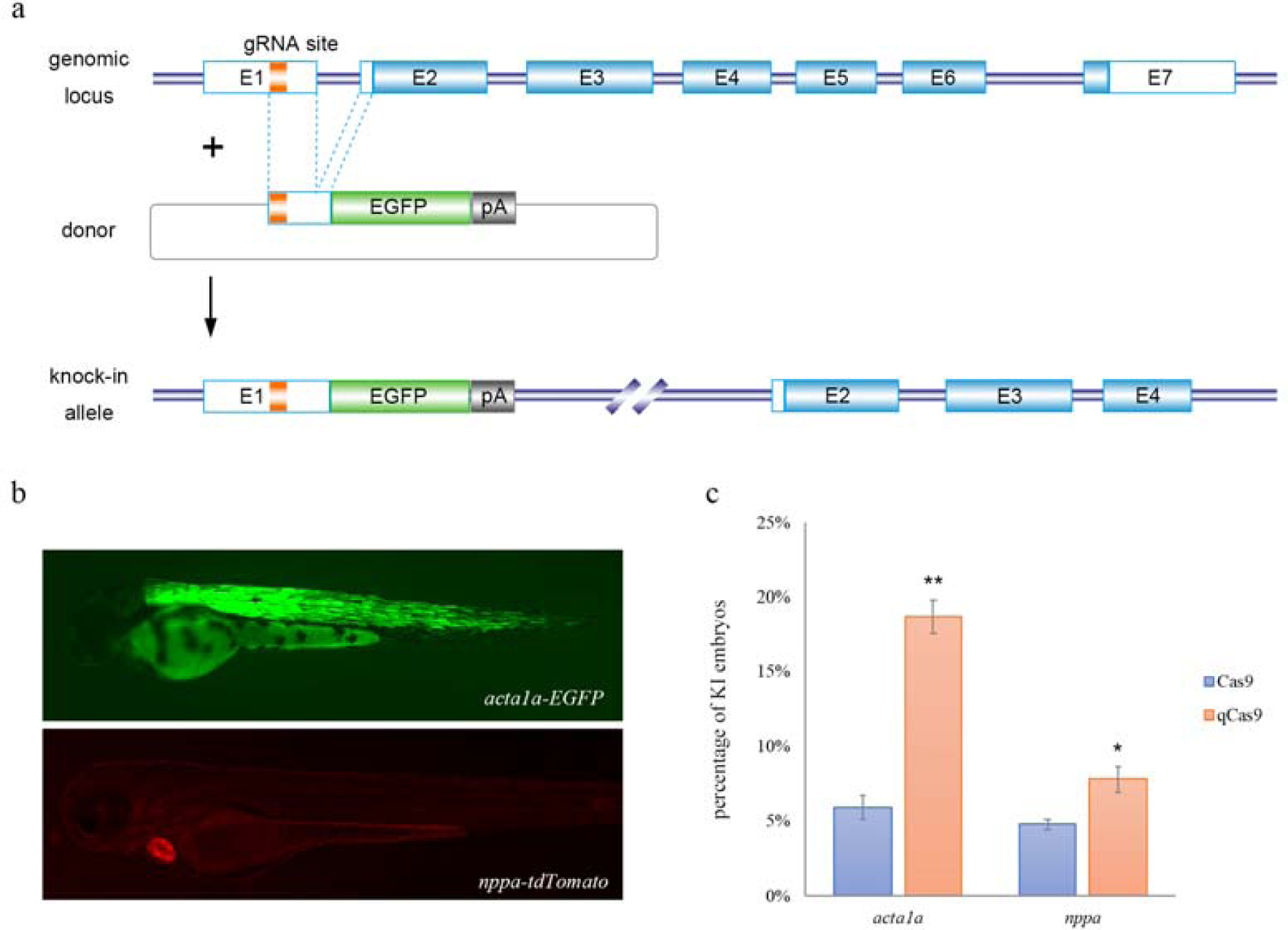
qCas9 increased targeted knock in efficiency at founder generation. (a) Schematic design of knock in donor and expected outcome. (b) Fluorescent images of positive knock in embryos with EGFP in trunk muscle (upper panel) and tdTomato in heart (lower panel). (c) Comparison of *acta1a*-EGFP positive knock in embryos and *nppa*-tdTomato positive knock in embryos after co-injection of respective knock in donor, gRNA and qCas9 or Cas9 mRNA. qCas9 significantly improved knock in efficiency from 5.9% (Cas9 injected, N=286) to 18.7% (qCas9 injected, N=346) for *acta1a*E1S1, and from 4.7% (Cas9 injected, N=336) to 7.8% (qCas9 injected, N=418) for *nppa*E1S2. Two-tailed unpaired Student’s t-test was used for statistical analysis with *, **, and *** represent *p* value < 0.05, 0.01 and 0.001, respectively.

### qCas9 led to higher germline transmission rate

To test whether the use of qCas9 could lead to higher germline transmission rate, founders of *smad5*E3S4 and *gch2*E1S1 were examined for germline transmission efficiencies. In Cas9/*smad5*E3S4 gRNA injected founders, only 1 of 12 showed germline transmission with 16.7% F_1_ offspring (4 of 24) carried mutant alleles. In contrast, 5 of 8 qCas9/*smad5*E3S4 gRNA injected founders showed germline transmission with the ratio of positive F_1_ offspring ranging from 33.3% to 83.3%. Similarly, in Cas9/*gch2*E1S1 gRNA injected founders, 9 of 11 showed germline transmission and the ratio of positive F_1_ offspring ranged from 25.0% to 75.0%, while 100% (6 of 6) qCas9/*gch2*E1S1 gRNA injected founders showed germline transmission with the ratio of positive F_1_ offspring ranging from 75.0% to 93.8% (**Table 1**). In sum, qCas9 can dramatically increase germline transmission rate, which holds great potential for facilitating founder screen process that are typically labor intensive.

**Table 1.**
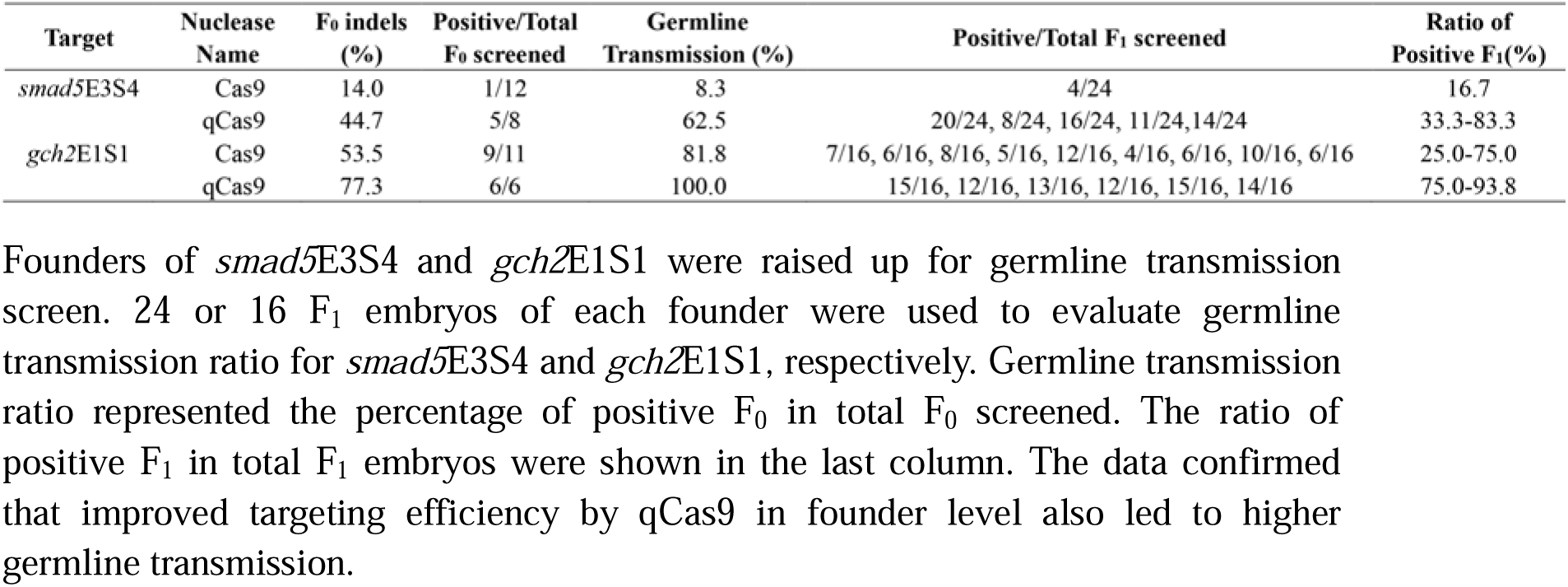
Germline transmission screen.

### qCas9 reduced off-target effect in HEK293T cell line

To explore whether qCas9 can be applied to mammalian systems, we fused the PEST sequence to the C terminal of human-codon-optimized Cas9 (quick hCas9, qhCas9) and compared targeting efficiency between qhCas9 and hCas9. Our results showed that only marginal improvements could be observed when HEK293T cells were transfected with qhCas9 compared to that of hCas9 together with gRNAs targeting five endogenous loci (**Supplementary Fig. 5**, target sites information in Supplementary Table 1). To determine whether the fast turnover rate of qhCas9 could help reduce off-target effect, two VEGFA target sites with pronounced off-target genome editing reported previously were tested in HEK293T cell line^28^. The potential off-target sites were amplified and subjected to next generation sequencing two days after transfection. The results revealed significant reduction in off-target editing when qhCas9 was used (**Figure 6**). In addition, the off-target editing of qCas9 in zebrafish was also tested. Potential off-target sites for *smad5*E3S4 (**Supplementary Table 2a**) and *tyr*E1S1 (**Supplementary Table 2b**) were predicted using CasOT program^47^ and analyzed using next generation sequencing. Our results showed no off-target editing was detected at 21 potential off-target sites of *smad5*E3S4 and 24 potential off-target sites of *tyr*E1S1(**Supplementary Fig. 6**). Since no off-target editing of Cas9 has been reported in zebrafish yet, comparative off-target analyses of more target loci are warranted for a conclusion of safer editing with qCas9. Taken together, the data demonstrates that qhCas9 can significantly reduce off-target editing and marginally improve on-target editing in HEK293T cell line.

**Figure 6.**
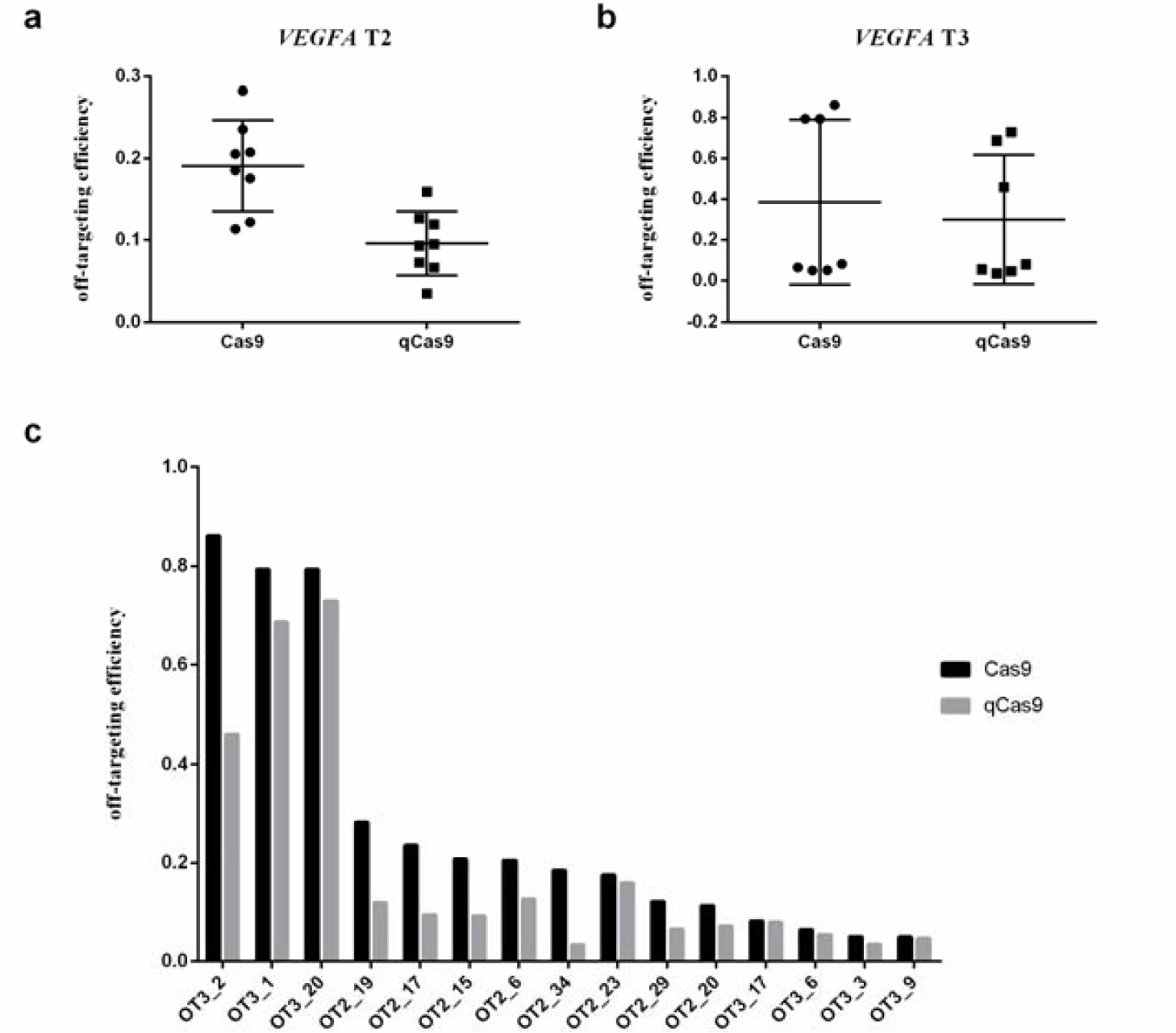
qCas9 significantly reduced off-target editing in HEK293T cell line. (a) Next generation sequencing results of editing efficiencies of potential off-target sites for VEGFA target site 2 (T2). (b) Next generation sequencing results of editing efficiencies of potential off-target sites for VEGFA target site 3 (T3). (c) Pairwise comparison of targeting efficiencies at potential off-target sites of VEGFA T2 and T3. qCas9 reduced targeting efficiencies at almost all sites with detectable off-target editing.

## Discussion

Here we report a simple but powerful strategy to improve targeting efficiency in zebrafish using the newly developed Cas9-PEST/gRNA system. The fusion of Cas9 and PEST sequence, which we term ‘quick Cas9, qCas9’, dramatically increases both Knock-out and Knock-in efficiencies at multiple endogenous genomic loci, leading to increased mutagenesis efficiency and higher germline transmission rate. Moreover, fusion of PEST sequence to hCas9 significantly reduces off-target effect in HEK293T cell line.

The fertilized egg of zebrafish divides extremely fast during the first couple of hours of development, and there are only about 4 germ cells at the end of 1000-cell stage^48^. Therefore, in order to achieve high targeting efficiency and subsequent germline transmission in zebrafish, once injected, Cas9/gRNA complex needs to work as fast as possible within a short time window. To address this, several strategies have been developed. Injection of Cas9 protein instead of mRNA saved the time used for Cas9 mRNA translation^23,49^, however, the requirement of expertise in purifying highly effective Cas9 protein in house and the high-cost of commercialized Cas9 protein may limit its use. Xie *et al* provided an alternative method based on microinjection of Cas9/gRNA into oocyte to improve targeting efficiency and germline transmission rate given that the Cas9 mRNA was antecedently expressed prior to fertilization^24^. However, this method requires sophisticated oocyte storage and micromanipulation procedures, which is not accessible to most of labs. A previous study also showed that adding ubiquitin-targeting signal to Cas9 could shorten its half-life and reduce mosaicism in non-human primate embryos while the overall targeting efficiency is not improved^50^. In this study, a short PEST sequence (40 amino acids) was added to Cas9, which resulted in significantly boosted targeting efficiency. The ability of PEST sequence in accelerating EGFP mRNA translation and protein degradation suggests the improvement in targeting efficiency of Cas9-PEST is in part due to faster Cas9-PEST protein turnover. It is worth noting that genome editing by qCas9 could be detected as early as 3 hpf, 1 hour earlier than the Cas9 protein as previously reported^23^, which may be attributed to the low targeting efficiency (<5%) of the previous study. In sum, qCas9 is a simple yet powerful tool to increase targeting efficiency.

Co-injection of qCas9 mRNA/gRNA improves targeting efficiency at multiple sites, especially for those with efficiencies less than 60%, thereby saving time and resource in finding high efficiency target sites required for Knock-in experiment. It is also important to keep in mind that many factors, such as chromatin and epigenetic state, sgRNA secondary structure, the affinity between sgRNA and Cas9 may all contribute to the variabilities in targeting efficiency at different genomic loci. While qCas9 increases targeting efficiency of one inactive target site to detectable level (<2%), the other three inactive sites examined in this study remain inactive. This suggests that qCas9 does not directly affect the chromatin and epigenetic state of each target locus. As for those sites with Cas9 efficiencies over 80%, there is little room for qCas9 to improve.

The improvement of targeting efficiency for qCas9 could be observed at low dosage (200pg, Figure 3d-3f). Dosage reduction translates into lower toxicity, which is desirable for experiments that need microinjection of multiple components, such as multiplex genome targeting and knock in experiments. In the meantime, the boost in targeting efficiency with qCas9 also leads to higher germline transmission rate, which can save time, labor and resource. Besides, higher targeting efficiency can lead to improved chance of obtaining biallelic loss-of-function mutations in founder generation. In the case of *tyr*, when compared to Cas9, although the targeting efficiency was not increased by qCas9 at 60 hpf, we observed more pronounced loss-of-pigmentation phenotype in qCas9 injected founders, which may result from higher targeting efficiencies of qCas9 at earlier developmental time.

Previous study showed that non-homologous single-strand DNA greatly stimulates Cas9-directed gene disruption, but the off-target editing is also increased^27^. In comparison, qCas9 increases on-target efficiency, while maintains similarly low level of off-target rate as Cas9 in zebrafish. Off-target editing may not be a big issue in zebrafish since breeding will eventually dilute out the off-target effects, however, for clinical and therapeutic applications where precise targeting is required due to safety concerns, the unwanted off-target editing poses a real problem. To check the off-target effects in human cells, we tested quick hCas9 system in HEK293T cell line. We only observed limited improvement on targeting efficiency compared to hCas9, which is not unexpected since the persistent expression of qhCas9 and hCas9 in cells could offset the fast turnover nature of PEST sequence and prolonged targeting window could also undercut the enhancement effect. Nonetheless, we observed dramatic reduction of off-target editing for several VEGFA off-target sites, which may be attributed to the fast turnover of qhCas9 protein that reduced undesirable targeting on unintended sites. Therefore, our qCas9 system offers a safer option in mammalian cell targeting with low cost especially for high-throughput manipulation.

## Materials and Methods

### Zebrafish husbandry and cultured cell line

All the zebrafish used in this study were maintained at standard condition in the zebrafish facility of Peking University. The wild type strain used in this study is Tübingen. HEK293T cell line was used for in vitro cell culture experiment.

### Plasmid construction

The zebrafish-codon-optimized Cas9 plasmid flanked by NLS at both N and C termini, the pGH-T7-zCas9, was described previously^25^. All the plasmids used in this study were constructed via seamless cloning method using ClonExpress MultiS One Step Cloning Kit (Vazyme Biotech, C113). For the construction of pGH-T7-zCas9-PEST fusion plasmid, the PEST domain sequence from mouse *Odc1* cDNA was cloned and inserted between the C termini of zCas9 and the C terminal NLS. The quick hCas9 vector was constructed by inserting PEST sequence into the C terminal of a human-codon-optimized Cas9. The coding sequence of EGFP was cloned into pCS2 vector to get pCS2-EGFP and PEST sequence was inserted at the C termini of EGFP CDS to get the pCS2-EGFP-PEST. The hEMX1-P2A-tdTomato knock-in donor plasmid was constructed by sequentially inserting a human EMX1 gRNA target site^51^, a 200 bp protection sequences, P2A sequence and tdTomato CDS into the pTST3 vector backbone. The *acta1a*-EGFP knock-in donor was constructed by inserting acta1a sgRNA target site and partial of *acta1a* 5’ UTR sequence into the upstream of EGFP.

### RNA synthesis

For making the mRNA, the template DNA was linearized by XbaI digestion (for Cas9 and qCas9 mRNA) or NotI digestion (for EGFP and EGFP-PEST mRNA) and then purified by TIANquick Mini Purification Kit (TIANGEN Biotech). The capped Cas9/qCas9 and EGFP/EGFP-PEST mRNA were synthesized by mMESSAGE mMACHINE T7 or SP6 kit (Invitrogen), respectively. The in vitro transcribed mRNA was purified by LiCl precipitation. The guide RNAs were synthesized as previously described^10^. The primers for generating DNA template of guide RNAs were designed manually by add a T7 promoter sequence to the 5’-upstream of gRNA sequence. The gRNAs were in vitro transcribed by T7 RNA Polymerase (Takara) and purified by LiCl precipitation or ethanol precipitation.

The detail method of LiCl precipitation: The in vitro transcription reaction (20 μL) took place at 37□ for 2 hours, upon completion, 1μL of DNase I was added for DNA template removal at 37□ for 20 min. 30 μL of LiCl and 30 μL of RNase-free water were added to the reaction tube and mixed up, the reaction tube was stayed at −20□ for at least 2 hours. Centrifuging at 13,000×g at 4□ for 20 min and discard the supernatant. Washing the RNA pellet with 500 μL 70% ethanol, centrifuge at 13,000×g at 4□ for 5 min and discard the supernatant. The RNA pellet was air-dried for 5 min at room temperature, dissolved in RNase-free water and quantified by Nanodrop (Thermo Fisher).

### Microinjection

Except for the dose-dependent experiment, 400 pg Cas9 or qCas9 mRNA and 50 pg gRNA were co-injected into one-cell stage zebrafish embryos. For the dose-dependent experiment, 200 pg, 400 pg, 600 pg and 800 pg Cas9 or qCas9 mRNA were co-injected with 50 pg gRNA into one-cell stage zebrafish embryos. For knock in experiment, the mixture of 400 pg Cas9 or qCas9 mRNA, 50 pg *nppa*E1s2 gRNA, 30 pg hEMX1 gRNA and 10 pg knock in donor was co-injected into one-cell stage zebrafish embryos. The injection method and dosages minimally affected the survival and normal development of injected embryos with more than 75% embryos survived and developed normally.

### Mutagenesis efficiency quantification

The small insertions and deletions (indels) efficiency induced by Cas9/gRNA and qCas9/gRNA were quantified by a restriction-endonuclease-resistant restriction fragment length polymorphism (RFLP) assay as reported^17^ or T7E1 assay ^26^. For RFLP assay, injected embryos with normal morphology at 24 hpf (hours post-fertilization) were collected in groups (3 embryos per group). Each group was treated with 30 μL 50 mmol/L NaOH at 95°C for 15 min, vortexed, and then neutralized by adding 1/10 volume of 1 mol/L pH 8.0 Tris-HCl buffer. The resulting genomic DNA was either used immediately as a PCR template or stored at −20°C. A short genomic region (∼300bp-700bp) flanking the target site was PCR amplified and 2 μL of the amplicon was used for restriction enzyme digestion. The percentage of uncleaved band (potential targeted alleles) was quantified by Image J software. For each case, 4 to 6 groups of embryos were analyzed. The uncleaved bands were gel extracted and subjected to TA cloning and then sequenced by Sanger sequencing. For T7 Endonucleases I assay, the PCR amplicons were purified using Gel Extraction kit (TIANGEN). A total of 200 ng of the purified PCR amplicon was denatured and slowly reannealed to facilitate heteroduplex formation. The reannealing procedure consisted of a 5 min denaturing step at 95°C, followed by cooling to 85°C at −2 °C per second and further to 25°C at −0.1 °C per second. The reannealed amplicon was then digested with 10 units of T7 Endonuclease I (New England Biolabs) at 37 °C for 90min. The reaction was stopped by adding 1 μL of 0.5M EDTA. The band intensity was quantified using ImageJ software. All the target sites information, the primers used for PCR amplification and restriction enzymes used for RFLP assay are listed in **Supplementary Table S1**.

### Germline transmission screening

Founder fish (F_0_) of *smad5*E3S4 and *gch2*E1S1 were outcrossed with wild type zebrafish to get F_1_ embryos. For each F_0_, 8 groups (2 or 3 embryos per group) of F_1_ embryos were collected for genomic DNA extraction at 24 hpf. RFLP assay was used for germline transmission screening.

### Assessment of off-target effect

The online CRISPR/Cas9 off-target prediction program CasOT^47^ was used to predict potential off-target sites in the whole zebrafish genome based on Ensembl database (GRCz10). We set the parameters to allow any of the four types of PAM, at most two mismatches in the 12-nt seed region of the target site, and at most five mismatches in the 8-nt non-seed region. A pair of primers was designed to amplify the region (∼120 bp) of each selected site, the barcode ATC, GCA and CGT were added to the 5’-end of the primer to amplify the genome regions from the qCas9/gRNA-injected embryos, Cas9/gRNA-injected embryos or wild type embryos, respectively. The PCR products of these sites were purified and mixed together for deep sequencing (Illumina). After a quality test, the sequencing data were categorized by the barcode and primer sequences, and then analyzed with Perl scripts to calculate the ratio of sequences with indels in the regions of each potential off-target site. Similarly, off-target sites for VEGFA T2 and T3^28^ were amplified and subjected to deep sequencing (BGIseq-500).

### Microscopy

The 2 dpf zebrafish embryos were anesthetized with 0.03% Tricaine (Sigma-Aldrich), mounted in 3% methylcellulose and imaged using a Zeiss Stemi 2000 Stereo microscope. The fluorescent images of EGFP and EGFP-PEST mRNA injected embryos were obtained by Zeiss Axio Imager Z1 microscope equipped with a Zeiss AxioCam MRc5 digital camera.

### Statistical testing

Pairwise comparisons between Cas9 and qCas9 samples were made using Student’s t-test.

## Supporting information

Supplementary Figure 1 on-target efficiency comparison

Supplementary Figure 1 on-target efficiency comparison

Supplementary Figure 1 on-target efficiency comparison

Supplementary Figure 2 time-course targeting efficiency comparison

Supplementary Figure 2 time-course targeting efficiency comparison

Supplementary Figure 2 time-course targeting efficiency comparison

Supplementary Figure 3 qCas9 and Cas9 mRNA titration

Supplementary Figure 4 dosage-dependent targeting efficiency comparison

Supplementary Figure 4 dosage-dependent targeting efficiency comparison

Supplementary Figure 4 dosage-dependent targeting efficiency comparison

Supplementary Figure 5 on-target efficiency comparison in HEK293T cell line

Supplementary Figure 6 off-target editing of qCas9 in zebrafish

## Acknowledgements

This research was supported by Shenzhen Municipal Government of China (No. JCYJ20160531194327655),Guangdong Provincial Key Laboratory of Genome Read and Write (No. 2017B030301011), Guangdong Provincial Academician Workstation of BGI Synthetic Genomics (No. 2017B090904014), and Shenzhen Peacock Plan (No.. KQTD20150330171505310).

We gratefully thank Yuying Gao, Yan Shen, Yingdi Jia and Jingliang Chen for zebrafish facility maintenance and colleagues at BGI-Shenzhen for technical support and discussion.

## Data Availability

BGISeq-500 sequencing reads for this study have been deposited in the CNSA (https://db.cngb.org/cnsa/) of CNGBdb with accession code CNP0000410.

---

Supplementary Table 1

Supplementary Table 2a

Supplementary Table 2b

Supplementary Figure 1 on-target efficiency comparison

Supplementary Figure 2 time-course targeting efficiency comparison

Supplementary Figure 3 qCas9 and Cas9 mRNA titration

Supplementary Figure 4 dosage-dependent targeting efficiency comparison

Supplementary Figure 5 on-target efficiency comparison in HEK293T cell line

Supplementary Figure 6 off-target editing of qCas9 in zebrafish

## Reference

1 Terns, M. P. & Terns, R. M. CRISPR-based adaptive immune systems. Current opinion in microbiology 14, 321–327, doi:10.1016/j.mib.2011.03.005 (2011).

2 Wiedenheft, B., Sternberg, S. H. & Doudna, J. A. RNA-guided genetic silencing systems in bacteria and archaea. Nature 482, 331–338, doi:10.1038/nature10886 (2012).

3 Bhaya, D., Davison, M. & Barrangou, R. CRISPR-Cas systems in bacteria and archaea: versatile small RNAs for adaptive defense and regulation. Annual review of genetics 45, 273–297, doi:10.1146/annurev-genet-110410-132430 (2011).

4 Jinek, M. et al. A programmable dual-RNA-guided DNA endonuclease in adaptive bacterial immunity. Science 337, 816–821, doi:10.1126/science.1225829 (2012).

5 Cong, L. et al. Multiplex Genome Engineering Using CRISPR/Cas Systems. Science 339, 819–823, doi:10.1126/science.1231143 (2013).

6 Mali, P. et al. RNA-guided human genome engineering via Cas9. Science 339, 823–826, doi:10.1126/science.1232033 (2013).

7 Li, D. et al. Heritable gene targeting in the mouse and rat using a CRISPR-Cas system. Nature biotechnology 31, 681–683, doi:10.1038/nbt.2661 (2013).

8 Wang, H. et al. One-step generation of mice carrying mutations in multiple genes by CRISPR/Cas-mediated genome engineering. Cell 153, 910–918, doi:10.1016/j.cell.2013.04.025 (2013).

9 Hwang, W. Y. et al. Efficient genome editing in zebrafish using a CRISPR-Cas system. Nat Biotech 31, 227–229, doi:(2013).

10 Chang, N. et al. Genome editing with RNA-guided Cas9 nuclease in zebrafish embryos. Cell research 23, 465–472, doi:10.1038/cr.2013.45 (2013).

11 Gratz, S. J. et al. Genome engineering of Drosophila with the CRISPR RNA-guided Cas9 nuclease. Genetics 194, 1029–1035, doi:10.1534/genetics.113.152710 (2013).

12 Friedland, A. E. et al. Heritable genome editing in C. elegans via a CRISPR-Cas9 system. Nature methods 10, 741–743, doi:10.1038/nmeth.2532 (2013).

13 Barnes, D. E. Non-homologous end joining as a mechanism of DNA repair. Current biology : CB 11, R455–457 (2001).

14 van den Bosch, M., Lohman, P. H. & Pastink, A. DNA double-strand break repair by homologous recombination. Biological chemistry 383, 873–892, doi:10.1515/bc.2002.095 (2002).

15 Urnov, F. D. et al. Highly efficient endogenous human gene correction using designed zinc-finger nucleases. Nature 435, 646–651, doi:10.1038/nature03556 (2005).

16 Sander, J. D. et al. Targeted gene disruption in somatic zebrafish cells using engineered TALENs. Nat Biotech 29, 697–698, doi:10.1038/nbt.1934 (2011).

17 Huang, P. et al. Heritable gene targeting in zebrafish using customized TALENs. Nature biotechnology 29, 699–700, doi:10.1038/nbt.1939 (2011).

18 Xiao, A. et al. Chromosomal deletions and inversions mediated by TALENs and CRISPR/Cas in zebrafish. Nucleic Acids Res 41, e141, doi:10.1093/nar/gkt464 (2013).

19 Doench, J. G. et al. Rational design of highly active sgRNAs for CRISPR-Cas9-mediated gene inactivation. Nature biotechnology 32, 1262–1267, doi:10.1038/nbt.3026 (2014).

20 Xu, H. et al. Sequence determinants of improved CRISPR sgRNA design. Genome research 25, 1147–1157, doi:10.1101/gr.191452.115 (2015).

21 Montague, T. G., Cruz, J. M., Gagnon, J. A., Church, G. M. & Valen, E. CHOPCHOP: a CRISPR/Cas9 and TALEN web tool for genome editing. Nucleic Acids Res 42, W401–407, doi:10.1093/nar/gku410 (2014).

22 Varshney, G. K. et al. High-throughput gene targeting and phenotyping in zebrafish using CRISPR/Cas9. Genome research 25, 1030–1042, doi:10.1101/gr.186379.114 (2015).

23 Sung, Y. H. et al. Highly efficient gene knockout in mice and zebrafish with RNA-guided endonucleases. Genome research 24, 125–131, doi:10.1101/gr.163394.113 (2014).

24 Xie, S. L. et al. A novel technique based on in vitro oocyte injection to improve CRISPR/Cas9 gene editing in zebrafish. Scientific reports 6, 34555, doi:10.1038/srep34555 (2016).

25 Liu, D. et al. Efficient gene targeting in zebrafish mediated by a zebrafish-codon-optimized cas9 and evaluation of off-targeting effect. Journal of genetics and genomics = Yi chuan xue bao 41, 43–46, doi:10.1016/j.jgg.2013.11.004 (2014).

26 Jao, L. E., Wente, S. R. & Chen, W. Efficient multiplex biallelic zebrafish genome editing using a CRISPR nuclease system. Proceedings of the National Academy of Sciences of the United States of America 110, 13904–13909, doi:10.1073/pnas.1308335110 (2013).

27 Richardson, C. D., Ray, G. J., Bray, N. L. & Corn, J. E. Non-homologous DNA increases gene disruption efficiency by altering DNA repair outcomes. Nature communications 7, 12463, doi:10.1038/ncomms12463 (2016).

28 Fu, Y. et al. High-frequency off-target mutagenesis induced by CRISPR-Cas nucleases in human cells. Nature biotechnology 31, 822–826, doi:10.1038/nbt.2623 (2013).

29 Fu, Y., Sander, J. D., Reyon, D., Cascio, V. M. & Joung, J. K. Improving CRISPR-Cas nuclease specificity using truncated guide RNAs. Nature biotechnology 32, 279–284, doi:10.1038/nbt.2808 (2014).

30 Ran, F. A. et al. Double nicking by RNA-guided CRISPR Cas9 for enhanced genome editing specificity. Cell 154, 1380–1389, doi:10.1016/j.cell.2013.08.021 (2013).

31 Tsai, S. Q. et al. Dimeric CRISPR RNA-guided FokI nucleases for highly specific genome editing. Nature biotechnology 32, 569–576, doi:10.1038/nbt.2908 (2014).

32 Slaymaker, I. M. et al. Rationally engineered Cas9 nucleases with improved specificity. Science 351, 84–88, doi:10.1126/science.aad5227 (2016).

33 Kleinstiver, B. P. et al. High-fidelity CRISPR–Cas9 nucleases with no detectable genome-wide off-target effects. Nature 529, 490–495, doi:10.1038/nature16526 (2016).

34 Wright, A. V. et al. Rational design of a split-Cas9 enzyme complex. Proceedings of the National Academy of Sciences of the United States of America 112, 2984–2989, doi:10.1073/pnas.1501698112 (2015).

35 Davis, K. M., Pattanayak, V., Thompson, D. B., Zuris, J. A. & Liu, D. R. Small molecule–triggered Cas9 protein with improved genome-editing specificity. Nature chemical biology 11, 316–318, doi:10.1038/nchembio.1793 (2015).

36 Zetsche, B., Volz, S. E. & Zhang, F. A split-Cas9 architecture for inducible genome editing and transcription modulation. Nat Biotech 33, 139–142, doi:10.1038/nbt.3149 (2015).

37 Polstein, L. R. & Gersbach, C. A. A light-inducible CRISPR-Cas9 system for control of endogenous gene activation. Nature chemical biology 11, 198–200, doi:10.1038/nchembio.1753 (2015).

38 Yin, H. et al. Partial DNA-guided Cas9 enables genome editing with reduced off-target activity. Nature chemical biology 14, 311–316, doi:10.1038/nchembio.2559 (2018).

39 Rogers, S., Wells, R. & Rechsteiner, M. Amino acid sequences common to rapidly degraded proteins: the PEST hypothesis. Science 234, 364–368 (1986).

40 Janne, O. A., Kontula, K. K., Isomaa, V. V. & Bardin, C. W. Ornithine decarboxylase mRNA in mouse kidney: a low abundancy gene product regulated by androgens with rapid kinetics. Annals of the New York Academy of Sciences 438, 72–84 (1984).

41 Rechsteiner, M. & Rogers, S. W. PEST sequences and regulation by proteolysis. Trends in biochemical sciences 21, 267–271 (1996).

42 Rechsteiner, M. PEST sequences are signals for rapid intracellular proteolysis. Seminars in cell biology 1, 433–440 (1990).

43 Ghoda, L., van Daalen Wetters, T., Macrae, M., Ascherman, D. & Coffino, P. Prevention of rapid intracellular degradation of ODC by a carboxyl-terminal truncation. Science 243, 1493–1495 (1989).

44 Ghoda, L., Phillips, M. A., Bass, K. E., Wang, C. C. & Coffino, P. Trypanosome ornithine decarboxylase is stable because it lacks sequences found in the carboxyl terminus of the mouse enzyme which target the latter for intracellular degradation. J Biol Chem 265, 11823–11826 (1990).

45 Li, X. et al. Generation of destabilized green fluorescent protein as a transcription reporter. J Biol Chem 273, 34970–34975 (1998).

46 Page-McCaw, P. S. et al. Retinal network adaptation to bright light requires tyrosinase. Nature neuroscience 7, 1329–1336, doi:10.1038/nn1344 (2004).

47 Xiao, A. et al. CasOT: a genome-wide Cas9/gRNA off-target searching tool. Bioinformatics (Oxford, England) 30, 1180–1182, doi:10.1093/bioinformatics/btt764 (2014).

48 Raz, E. Primordial germ-cell development: the zebrafish perspective. Nature reviews. Genetics 4, 690–700, doi:10.1038/nrg1154 (2003).

49 Gagnon, J. A. et al. Efficient mutagenesis by Cas9 protein-mediated oligonucleotide insertion and large-scale assessment of single-guide RNAs. PLoS One 9, e98186, doi:10.1371/journal.pone.0098186 (2014).

50 Tu, Z. et al. Promoting Cas9 degradation reduces mosaic mutations in non-human primate embryos. Scientific reports 7, 42081, doi:10.1038/srep42081 (2017).

51 Lin, S., Staahl, B. T., Alla, R. K. & Doudna, J. A. Enhanced homology-directed human genome engineering by controlled timing of CRISPR/Cas9 delivery. eLife 3, e04766, doi:10.7554/eLife.04766 (2014).

