## Supplementary figures and images for "Enhance genome editing efficiency and specificity by a quick CRISPR/Cas9 system"

### Supplementary Figure 1 on-target efficiency comparison

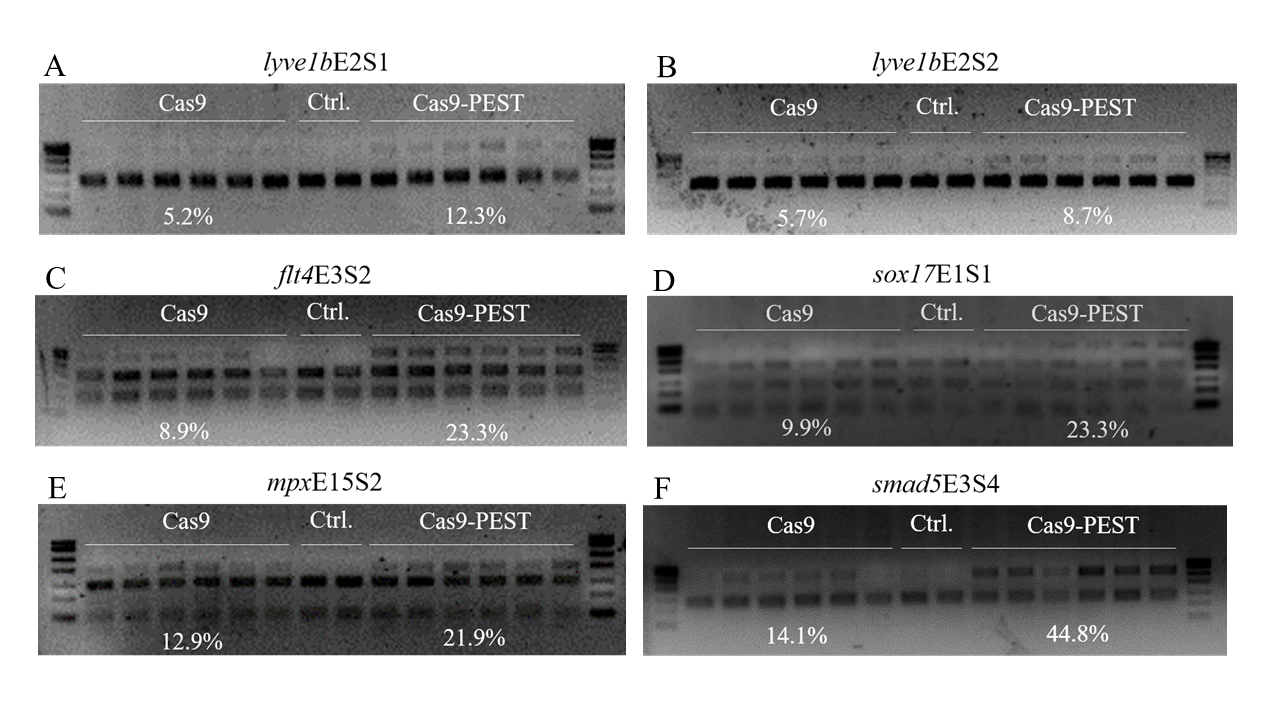

### Supplementary Figure 1 on-target efficiency comparison

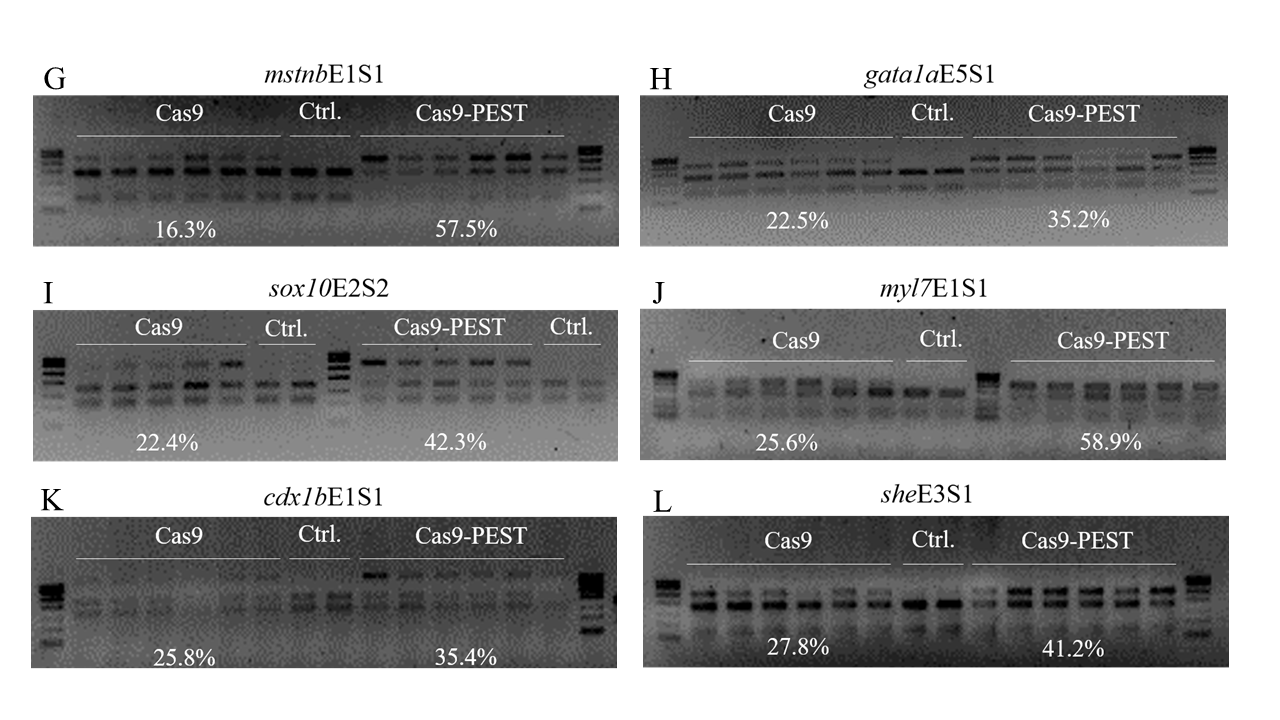

### Supplementary Figure 1 on-target efficiency comparison

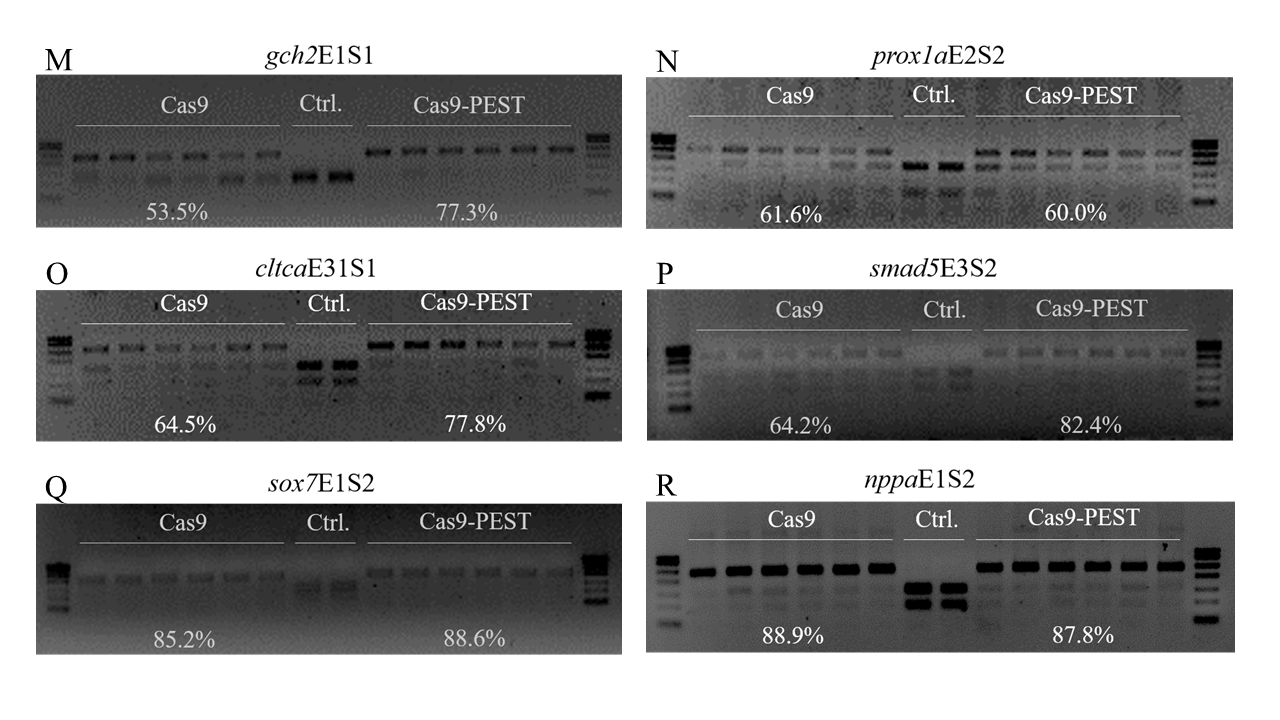

### Supplementary Figure 2 time-course targeting efficiency comparison

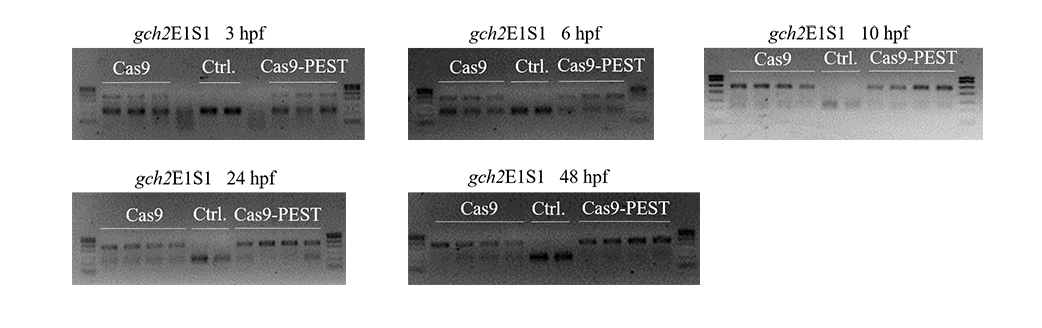

### Supplementary Figure 2 time-course targeting efficiency comparison

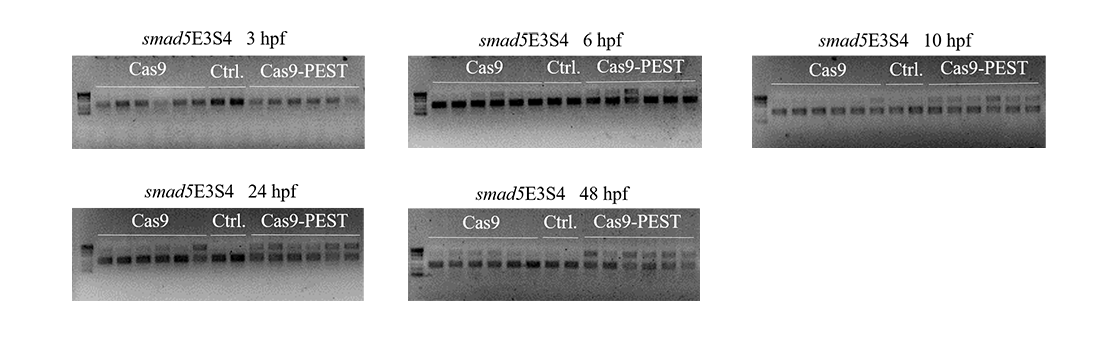

### Supplementary Figure 2 time-course targeting efficiency comparison

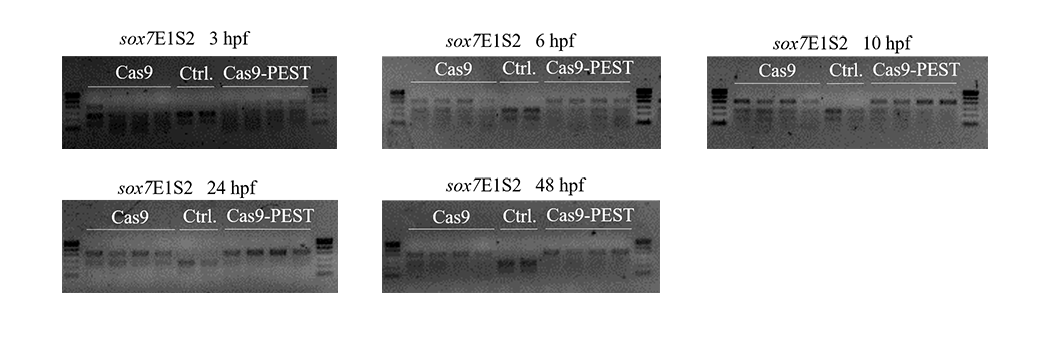

### Supplementary Figure 3 qCas9 and Cas9 mRNA titration

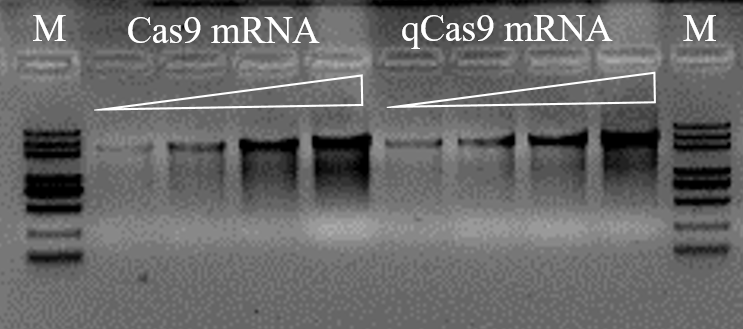

### Supplementary Figure 4 dosage-dependent targeting efficiency comparison

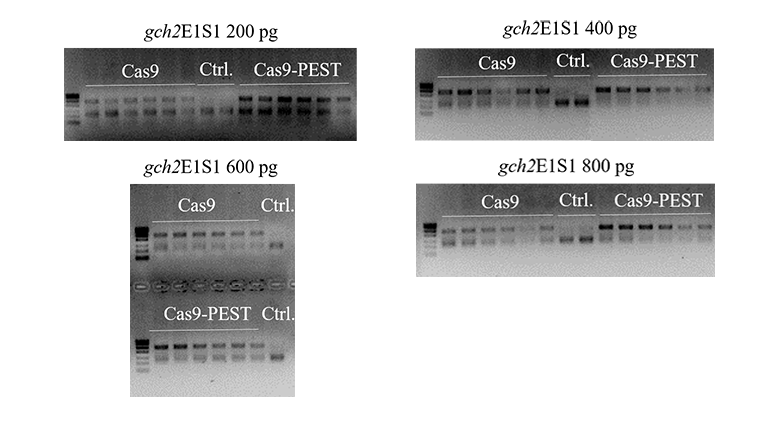

### Supplementary Figure 4 dosage-dependent targeting efficiency comparison

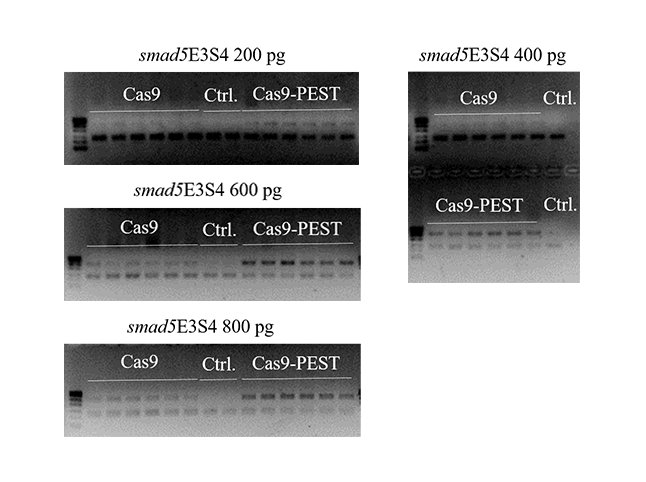

### Supplementary Figure 4 dosage-dependent targeting efficiency comparison

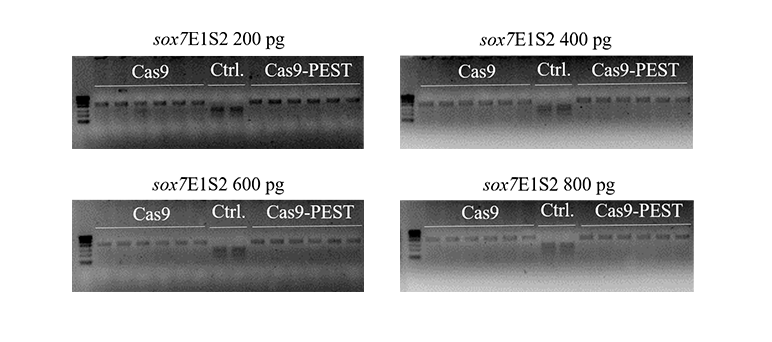

### Supplementary Figure 5 on-target efficiency comparison in HEK293T cell line

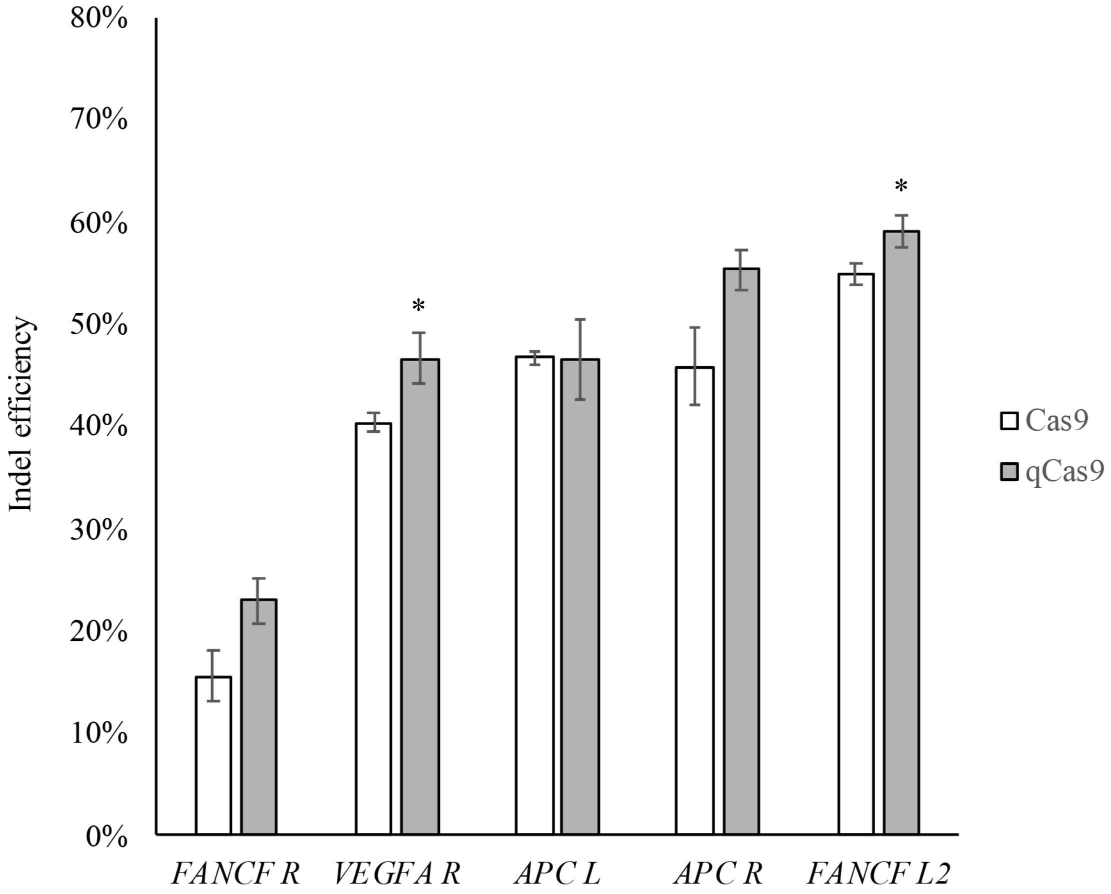

### Supplementary Figure 6 off-target editing of qCas9 in zebrafish

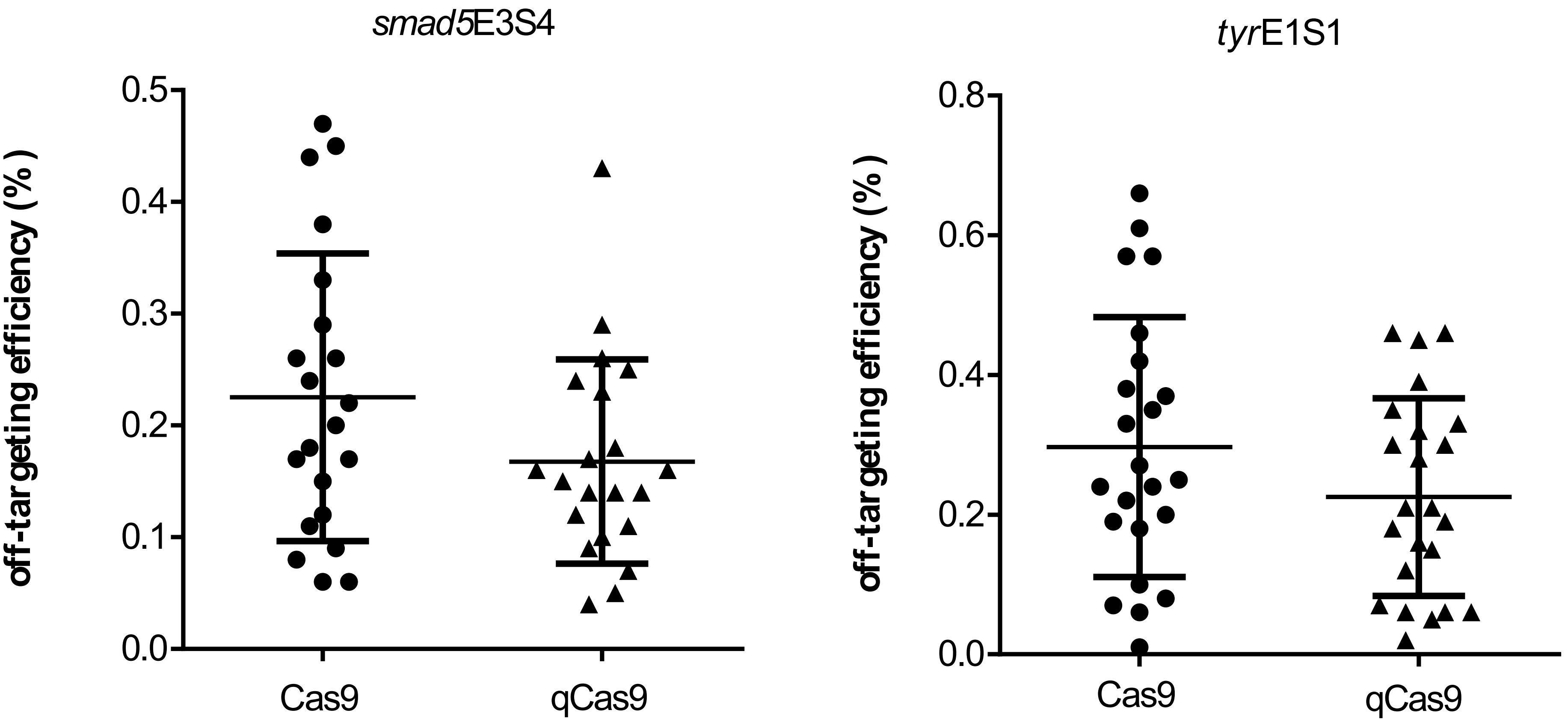
