## Supplementary Table 1 & 2 for "Enhance genome editing efficiency and specificity by a quick CRISPR/Cas9 system"

**Supplementary Information**

**Table S1. Cas9/gRNA target site information**

| Target name | Cas9/gRNA target site | Enzyme | Forward primer | Reverse primer | Amplicon size |
| --- | --- | --- | --- | --- | --- |
| *cdx1b*E1S1 | GGAGTGTCCAG**CTCGAG**CACAGG | XhoI | ACGTCACTGCTCCAACCTTT | GGTGTCAGAAACAGAATTTGGC | 720 bp |
| *cltca*E31S1 | GTGGTGGGACGGGCAC**ACTGG**GG | BsrI | TGTCACGTTGTACCAGCCG | GAGGGAGATCCATGGGGTTTG | 514 bp |
| *flt4*E3S2 | GCTGTTATTGAT**GGCACC**ACGG | BanI | TGTAGCACCTTGGTCCTGG | CGATTCGGTTGCACATTCC | 568 bp |
| *gata1a*E5S1 | GTAGTGTTGTAGT**ACTAGT**GTGG | SpeI | GCATTTAGTTCACCAGAAGCG | CCTGGGTTCAGAGAATACGC | 498 bp |
| *gch2*E1S1 | ACAGAGTATCTCT**GCCGCAATGG** | BglI | GAATCTGAGTCAGCTCCACG | CGCTTACCGTAGATGGTCTC | 453 bp |
| *gch2*E1S2 | GCGGTTACCTGC**TCTGGA**GGCGG | Hpy188III | GAATCTGAGTCAGCTCCACG | CGCTTACCGTAGATGGTCTC | 453 bp |
| *lyve1b*E2S1 | GATGCCATCACA**GCCAAAGAGG** | BglI | AATCATGGCTACAGTCTGCG | GTGCTTGACAGTCCTTTGGT | 579 bp |
| *lyve1b*E2S2 | ACAT**GCCTCTTTGGC**TGTGATGG | BglI | AATCATGGCTACAGTCTGCG | GTGCTTGACAGTCCTTTGGT | 579 bp |
| *mpx*E15S1 | GCAGGACCTC**CCGGACCCAGG**GG | BslI | AATGCTTTGTTGTTGTTTAATGAC | CAACTTTAGCAGTGGCAGGA | 388 bp |
| *mpx*E15S2 | GGAGGTCC**CCTGGGTCCGG**GAGG | BslI | AATGCTTTGTTGTTGTTTAATGAC | CAACTTTAGCAGTGGCAGGA | 388 bp |
| *mstnb*E1S1 | TCCAAACATCAG**CCGGGACGTGG** | BslI | ATGGAGATATAACGGCGCAC | ATGCGTAAAATTGCTGTGGC | 547 bp |
| *myl7*E1S1 | GGCTCTGGGTGT**CCATGTAGGGG** | BslI | GCACCTTTTACACACACAGC | TCACCATGTTCACTGTCTGC | 510 bp |
| *myl7*E2S1 | AAAAAGCCGCGG**CCAAGAGGGGG** | BslI | ACCCTCAACACTGAGTTCAA | TGGACATACACTGAATACAAACCA | 457 bp |
| *myl7*E2S2 | GTTTT**CCCCCTCTTGG**CCGCGG | BslI | ACCCTCAACACTGAGTTCAA | TGGACATACACTGAATACAAACCA | 457 bp |
| *nppa*E1S2 | CAGGACTGCTGCT**CCTGG**TTTGG | BstNI | CCAGCAAGCGTTATCACAGA | GTAAACATGCTGCCAATGACC | 471 bp |
| *nppa*E2S1 | TCGACAATAGG**AGATCT**AGCAGG | BglII | TATGAAGACAGCAACACCGT | ACACTCAATAGAGTCTTATTCGTT | 490 bp |
| *nppa*E2S2 | GTTTGCTTCGG**GTCGAC**AATAGG | SalI | TATGAAGACAGCAACACCGT | ACACTCAATAGAGTCTTATTCGTT | 490 bp |
| *prox1a*E2S2 | GGAAGTGGCCT**CTCGAG**TACCGG | XhoI | GCCTGACCATGACAGCACA | GCACGTTTGGCCCTTAGAT | 498 bp |
| *she*E3S1 | GGGACATCAT**AGATCT**GAGGGGG | BglII | TGTTCTGCAAAATTGCTCCACATGG | GGGGAGTTACCTGACAGGGCT | 492 bp |
| *smad5*E3S2 | GCGGCAGTAGATG**ACGTGT**GGG | AflIII | GGGCGATGAGGAGGAAAAAT | CCGGGGTAAAATATGGAGGC | 500 bp |
| *smad5*E3S4 | GGATACTCGCAAAC**CTCGAG**CGG | XhoI | GGGCGATGAGGAGGAAAAAT | CCGGGGTAAAATATGGAGGC | 500 bp |
| *sox7*E1S2 | GGCTCCGACACCTTGT**CCGCGG**G | SacII | TGCTCACTCAAACCCATCTTT | CAACGTTAAAATCTTACCAAGC | 387 bp |
| *sox10*E2S1 | GCGGTCAGT**CAGGTGCTG**AACGG | AlwNI | GTCGGAGGTGGAAATGAGTC | GGTAAAGCATCATTCGTGCG | 444 bp |
| *sox10*E2S2 | GAGTTCACGCG**CACGGGCATG**GG | MslI | GTCGGAGGTGGAAATGAGTC | GGTAAAGCATCATTCGTGCG | 444 bp |
| *sox17*E1S1 | CATGACTGAG**CTGCAG**CTGCTGG | PstI | GCTATTGGTGGCTGCTGAGT | TTGTGCAGCATGGGCATAGT | 678 bp |
| *tyr*E1S1 | GGACTGGAGGACTTCTGGGGAGG | T7E1 | GCGTCTCACTCTCCTCGACTCTTC | GTAGTTTCCGGCGCACTGGCAG | 315 bp |
| *EMX1* | GTCACCTCCAATGACTAGGGTGG | T7E1 | GCCATCCCCTTCTGTGAATGTTAGAC | GGAGATTGGAGACACGGAGAGCAG | 640 bp |
| APC L | CCAGAAGTACGAGC**GCCGCCCGG** | BglI | CTGTTCCCAGGTACTGTTGT | GAGAATGGAGGACCTGCAAAG | 435bp |
| APC R | TGGCAGGTGAGTGAGG**CTGCAG**G | PstI | CTGTTCCCAGGTACTGTTGT | GAGAATGGAGGACCTGCAAAG | 435bp |
| FANCL R | GGAATCCCTT**CTGCAG**CACCTGG | PstI | CATCTGCTCTCCCTCCACTA | CTGGAAGTTCGCTAATCCCG | 481bp |
| FANCL L2 | TTCCGCTTTCACCTTG**GAGACG**G | BsmBI | CATCTGCTCTCCCTCCACTA | CTGGAAGTTCGCTAATCCCG | 481bp |
| VEGFA R | TCCCTCTTTAGCCAGAG**CCGG**GG | MspI | CGTTCTCAGCTCCACAAACT | GAATCCTGGAGTGACCCCT | 489bp |

The sequence information of Cas9/gRNA target sites, corresponding primers used for the PCR amplification and amplicon size. Note that PAM sequences are marked in red and restriction sites are marked bold.

**Table S2a. *smad5*E3S4 off-target sites information**

| **Site name** | **Target site sequence** | **Genomic location** | **Indel (%)** | | |
| --- | --- | --- | --- | --- | --- |
|  |  |  | **Control** | **Cas9** | **qCas9** |
| *smad5*E3S4 | GGATACTCGCAAACCTCGAGCGG | *smad5* exon3 | 0.59 | 3.62 | 14.97 |
| *smad5* OT1 | AGGTGATCGCAAACCTCGAGAGG | *lck* exon13 | 0.05 | 0.08 | 0.12 |
| *smad5* OT2 | ATATCCTAGCAAACCTCGAGAGG | *tbc1d12b* intron5 | 0.10 | 0.12 | 0.09 |
| *smad5* OT3 | GGATGATCGCAAAACTCGGGTGG | intergenic | 0.20 | 0.33 | 0.26 |
| *smad5* OT4 | AGAGACAGGCAAAGCTCGAGCGG | *mrpl17* exon1 | 0.10 | 0.20 | 0.05 |
| *smad5* OT5 | TGATTTTAGCAAACCACGAGAGG | intergenic | 0.15 | 0.11 | 0.15 |
| *smad5* OT6 | TCTTTCTCGCAAACCTCAAGTGG | intergenic | 0.33 | 0.47 | 0.43 |
| *smad5* OT7 | GCATGCTGGGAAATCTCGAGAGG | *myo7ab* exon43 | 0.23 | 0.18 | 0.29 |
| *smad5* OT8 | AGATGCGCGCAAACCTAGAAAGG | *ptprdb* intron18 | 0.23 | 0.26 | 0.16 |
| *smad5* OT9 | CGGTTCTCGGAAACATCGAGTGG | intergenic | 0.03 | 0.06 | 0.04 |
| *smad5* OT10 | AGATTTTCTCAAACATCGAGGGG | *adgrb3* intron14 | 0.27 | 0.17 | 0.17 |
| *smad5* OT11 | TAACACTCGCAAAACACGAGAGG | *EML5* intron17 | 0.20 | 0.22 | 0.10 |
| *smad5* OT12 | ACATACACACAAACCTCGATAGG | intergenic | 0.03 | 0.06 | 0.07 |
| *smad5* OT13 | GGATTCCTGCAACCCTGGAGGGG | *cacna2d4a* intron6 | 0.22 | 0.44 | 0.24 |
| *smad5* OT14 | GCTCAAGCGCAAACTTCGAGAGG | *ralbp1* exon5 | 0.22 | 0.24 | 0.25 |
| *smad5* OT15 | GTAACATTGCAAACCTCAAGTGG | intergenic | 0.14 | 0.38 | 0.18 |
| *smad5* OT16 | TTTTCCTGGCAAGCCTCGAGAGG | *bcas3* intron22 | 0.10 | 0.09 | 0.16 |
| *smad5* OT17 | AGCTCCATGCAAACCTTGAGAGG | *creb5b* intron5 | 0.30 | 0.29 | 0.14 |
| *smad5* OT18 | AATCACTGGCAAACCTTGAGCGG | intergenic | 0.32 | 0.26 | 0.23 |
| *smad5* OT19 | AAATAAAAGCCAACCTCGAGCGG | *sar1b* intron1 | 0.17 | 0.15 | 0.11 |
| *smad5* OT20 | AGCAATTAGCAAACCTCCAGAGG | *camk2b1* intron4 | 0.38 | 0.45 | 0.14 |
| *smad5* OT21 | TTCCGCTCGCAAACTTCGAGGGG | intergenic | 0.18 | 0.17 | 0.14 |

**Table S2b. *tyr*E1S1 off-target sites information**

| **Site name** | **Target site sequence** | **Genomic location** | **Indel (%)** | | |
| --- | --- | --- | --- | --- | --- |
|  |  |  | **Control** | **Cas9** | **qCas9** |
| *tyr* E1S1 | GGACTGGAGGACTTCTGGGGAGG | *tyr* exon1 | 0.61 | 87.33 | 85.43 |
| *tyr* OT1 | TGACTGGAGGCCTTTTGGGGGGG | intergenic | 0.01 | 0.01 | 0.02 |
| *tyr* OT2 | GAGCTGTAGGACTTCTGGGTGGG | *tenm3* exon1 | 0.34 | 0.46 | 0.45 |
| *tyr* OT3 | CATCTGGAGGACTTCTGAGGCGG | *gbe1b* intron10 | **0.70** | **45.78** | **46.92** |
| *tyr* OT4 | AGATTGAAGGACTTCTGGGCTGG | *DST* exon2 | 0.40 | 0.22 | 0.46 |
| *tyr* OT5 | TGACTGATGGACTTCTGAGGCGG | *rab3ip* exon3 | 0.29 | 0.07 | 0.28 |
| *tyr* OT6 | TGACTGCAGGACTGCTGAGGTGG | intergenic | 0.10 | 0.24 | 0.16 |
| *tyr* OT7 | TAACTGGAGGTCTTTTGGGGGGG | intergenic | 0.11 | 0.24 | 0.07 |
| *tyr* OT8 | GGAGCGGAGGGCTTCAGGGGAGG | intergenic | 0.19 | 0.27 | 0.19 |
| *tyr* OT9 | AAACAGTAGGACATCTGGGGAGG | *plxna4* intron2 | 0.31 | 0.42 | 0.30 |
| *tyr* OT10 | GTGCTGCTGGACTCCTGGGGTGG | intergenic | 0.05 | 0.18 | 0.06 |
| *tyr* OT11 | TCAGTGGAGGATTTTTGGGGCGG | *pla2r1* exon2 | 0.30 | 0.57 | 0.35 |
| *tyr* OT12 | TTACTGGGGGAGATCTGGGGAGG | *bcas3* exon17 | 0.33 | 0.61 | 0.18 |
| *tyr* OT13 | GAACGGGCGGTCTTCTGCGGTGG | *trim71* exon1 | 0.32 | 0.35 | 0.39 |
| *tyr* OT14 | GTATTTGAGGATTTCTGGTGCGG | *rps6ka5* exon14 | 0.32 | 0.57 | 0.30 |
| *tyr* OT15 | ATACTCGAGGACCTCTGGGCAGG | *zfyve9a* exon2 | 0.31 | 0.33 | 0.32 |
| *tyr* OT16 | GGACGTGCGGACTTCAGAGGTGG | *taok1b* exon1 | 0.06 | 0.08 | 0.06 |
| *tyr* OT17 | GAAGTGCAGAACTTCTGGCGGGG | *TXK* exon12 | 0.41 | 0.66 | 0.46 |
| *tyr* OT18 | GGAGTGCTGGACCTATGGGGAGG | *si:dkeyp-89c11.3* exon6 | 0.29 | 0.19 | 0.21 |
| *tyr* OT19 | TGAGAGGAGGGTTTCTGGGGAGG | *card9* exon11 | 0.09 | 0.10 | 0.06 |
| *tyr* OT20 | CACCTGGAAGAGTTCTGGGGAGG | intergenic | 0.01 | 0.06 | 0.05 |
| *tyr* OT21 | GCCCAGGAGGGCTTCTGAGGAGG | *ptgir* intron2 | 0.37 | 0.38 | 0.33 |
| *tyr* OT22 | GGACATCAGGCCTTCAGGGGAGG | intergenic | 0.22 | 0.20 | 0.15 |
| *tyr* OT23 | TGACAGAAGGACGTCAGGGGAGG | intergenic | 0.18 | 0.25 | 0.12 |
| *tyr* OT24 | GGAAAGCATGACTTCTGTGGCGG | *arih1l* intron12 | 0.28 | 0.37 | 0.21 |
